# Adhesion G protein-coupled receptor ADGRG1 promotes protective microglial response in Alzheimer’s disease

**DOI:** 10.1101/2024.10.15.618329

**Authors:** Beika Zhu, Andi Wangzhou, Diankun Yu, Tao Li, Rachael Schmidt, Stacy L. De Florencio, Lauren Chao, Yonatan Perez, Lea T. Grinberg, Salvatore Spina, Richard M. Ransohoff, Arnold R. Kriegstein, William W. Seeley, Tomasz Nowakowski, Xianhua Piao

**Affiliations:** Weill Institute for Neuroscience, University of California, San Francisco, San Francisco, CA 94158, USA; Eli and Edythe Broad Center of Regeneration Medicine and Stem Cell Research, University of California, San Francisco, San Francisco, CA 94143, USA; Department of Neurology, University of California, San Francisco, San Francisco, CA 94143, USA; Department of Pathology, University of California, San Francisco, San Francisco, CA 94143, USA; Memory and Aging Center, University of California, San Francisco, San Francisco, CA 94158, USA; Third Rock Ventures, Boston, MA 02215, USA; Department of Anatomy, University of California, San Francisco, San Francisco, CA 94143, USA; Department of Psychiatry, University of California, San Francisco, San Francisco, CA 94143, USA; Department of Neurological Surgery, University of California, San Francisco, San Francisco, CA 94143, USA; Kavli Institute for Fundamental Neuroscience, University of California, San Francisco, San Francisco, CA 94143, USA; Division of Neonatology, Department of Pediatrics, University of California at San Francisco, San Francisco, CA 94158, USA; Newborn Brain Research Institute, University of California at San Francisco, San Francisco, CA 94158, USA

## Abstract

Germline genetic architecture of Alzheimer’s disease (AD) indicates microglial mechanisms of disease susceptibility and outcomes. However, the mechanisms that enable microglia to mediate protective responses to AD pathology remain elusive. *Adgrg1* is specifically expressed in yolk-sac-derived microglia. This study reveals the role of yolk-sac-derived microglia in AD pathology, highlighting the function of ADGRG1 in modulating microglial protective responses to amyloid deposition. Utilizing both constitutive and inducible microglial *Adgrg1* knockout 5xFAD models, we demonstrate that *Adgrg1* deficiency leads to increased amyloid deposition, exacerbated neuropathology, and accelerated cognitive impairment. Transcriptomic analyses reveal a distinct microglial state characterized by downregulated genes associated with homeostasis, phagocytosis, and lysosomal functions. Functional assays in mouse models and human embryonic stem cells-derived microglia support that microglial ADGRG1 is required for efficient Aβ phagocytosis. Together, these results uncover a GPCR-dependent microglial response to Aβ, pointing towards potential therapeutic strategies to alleviate disease progression by enhancing microglial functional competence.

## Introduction

Alzheimer’s disease (AD) is a prevalent form of late-life dementia that progresses relentlessly and is ultimately fatal. Pathologically, AD is characterized by the accumulation of extracellular amyloid beta (Aβ) senile plaques and intracellular neurofibrillary tangles composed of hyperphosphorylated tau.^1–3^ These proteinaceous aggregates cause extensive neuritic dystrophy, as well as synaptic and neuronal loss, cumulatively leading to a gradual decline in memory and cognitive functions.^1–4^ Microglia, the resident innate immune cells of the central nervous system (CNS), play important roles in responding to these pathological changes. Genome-wide association studies (GWAS) have identified numerous loci associated with AD, with several genes being highly expressed in microglia. For instance, *BIN1*, a major AD GWAS hit, contains a risk single nucleotide polymorphism (SNP) located in a microglia-specific enhancer.^5^ Microglia are known for their remarkable plasticity, dynamically adapting their transcriptomic and functional states in response to various environmental cues.^6–8^ The mechanisms guiding these state transitions, particularly those fostering protective responses in AD, are not fully understood. Importantly, the properties and states of microglia are determined by their ontogeny and the CNS environment. Although transplanted macrophages or hematopoietic stem cells from bone-marrow or blood can express microglial genes in the brain, only those of yolk-sac origin fully attain microglial identity.^9,10^ *Adgrg1* is one of the few genes exclusively expressed in yolk-sac-derived microglia, but not in other microglia-like cells.^9^ ADGRG1 is an adhesion G protein-coupled receptor (aGPCR) that plays critical roles in brain development. Loss-of-function mutations in *ADGRG1* result in severe human brain malformations,^11,12^ and deletion of microglial *Adgrg1* impairs synaptic pruning and interneuron development in the early postnatal mouse brain.^13,14^ However, the function of microglial ADGRG1 in AD pathology remains uncharacterized.

In this study, we extend our understanding of the roles of yolk-sac derived microglia in AD pathology by cell-specific deletion of microglial *Adgrg1* in the 5xFAD mouse model. We demonstrate that targeted deletion of *Adgrg1* in microglia leads to higher plaque deposition, increased neuronal and synaptic loss, and deficits in spatial memory in 5xFAD mice. Through single-nucleus RNA sequencing (snRNA-seq), we identify unique microglial transcriptomic changes induced by *Adgrg1* deletion in 5xFAD mice, with significantly reduced expression of genes related to homeostasis, phagocytosis, and lysosomal functions. Additionally, our study shows that deleting microglial *ADGRG1* leads to impaired associations between microglia and amyloid plaques, as well as deficiencies in microglial phagocytic activity in both primary mouse microglia and human embryonic stem cells (hESC)-derived microglia. Collectively, these findings provide insights into the role of yolk-sac-derived microglia in AD pathology, uncovering an underrecognized role of ADGRG1 in mediating protective microglial responses during AD progression.

## Results

### Microglial *Adgrg1* deletion exacerbates Aβ plaque load and impairs microglia-plaque association

To elucidate the role of microglial ADGRG1 in AD pathology, we first employed *Cx3cr1^Cre(Jung)^* transgenic mice,^15^ based on our past success with this Cre driver.^13,14^ We verified the specificity of Cre-Lox recombination in microglia using the *Cx3cr1^Cre^* driver by crossing it with a *Rosa-GFP^fl^* reporter line, confirming that GFP expression was exclusively localized to microglia.^14^ Although *Cx3cr1^Cre/+^* induces recombination in all myeloid cells, *Adgrg1* is selectively expressed in microglia, but not in other tissue mononuclear phagocytes.^9,16^ We chose 5xFAD^17^ as our AD mouse model based on the following two reasons: (1) their aggressive phenotype allows us to screen the most profound functions of microglial ADGRG1 in a relatively short time, and (2) there is a large amount of transcriptomic data available in the public domain that can be mined and compared. Specifically, this allows us to take advantage of the data generated by Marco Colonna’s group on *Trem2* deletion in 5xFAD mice to detect ADGRG1-specific functions in mediating microglial response to Aβ. We crossed *Adgrg1^fl/fl^;Cx3cr1^Cre/+^*with 5xFAD mice to generate conditional knockout mice *5xFAD;Adgrg1^fl/fl^;Cx3cr1^Cre/+^*(5xFAD-cKO) and age-matched controls, *5xFAD;Adgrg1^+/+^;Cx3cr1^Cre/+^* mice (5xFAD-Con) (**Figure 1a**). We confirmed specific deletion of *Adgrg1* in microglia (**Figures S1a-c**). We first examined Aβ plaque accumulation via immunofluorescence for MOAB2^18^ and observed a significant increase in Aβ deposition in the cortex and hippocampal CA1 region of 6-month-old 5xFAD-cKO mice compared to 5xFAD-Con mice (**Figures 1b-d**, **S1d-e**). Further analysis using Thioflavin S (ThioS) and MOAB2 double staining, where ThioS labels the compact cores of fibrillar and protofibrillar Aβ,^19,20^ revealed that plaques in 5xFAD-cKO mice were more diffuse compared to those in 5xFAD-Con mice (**Figures 1e-f**).

**Figure 1.**
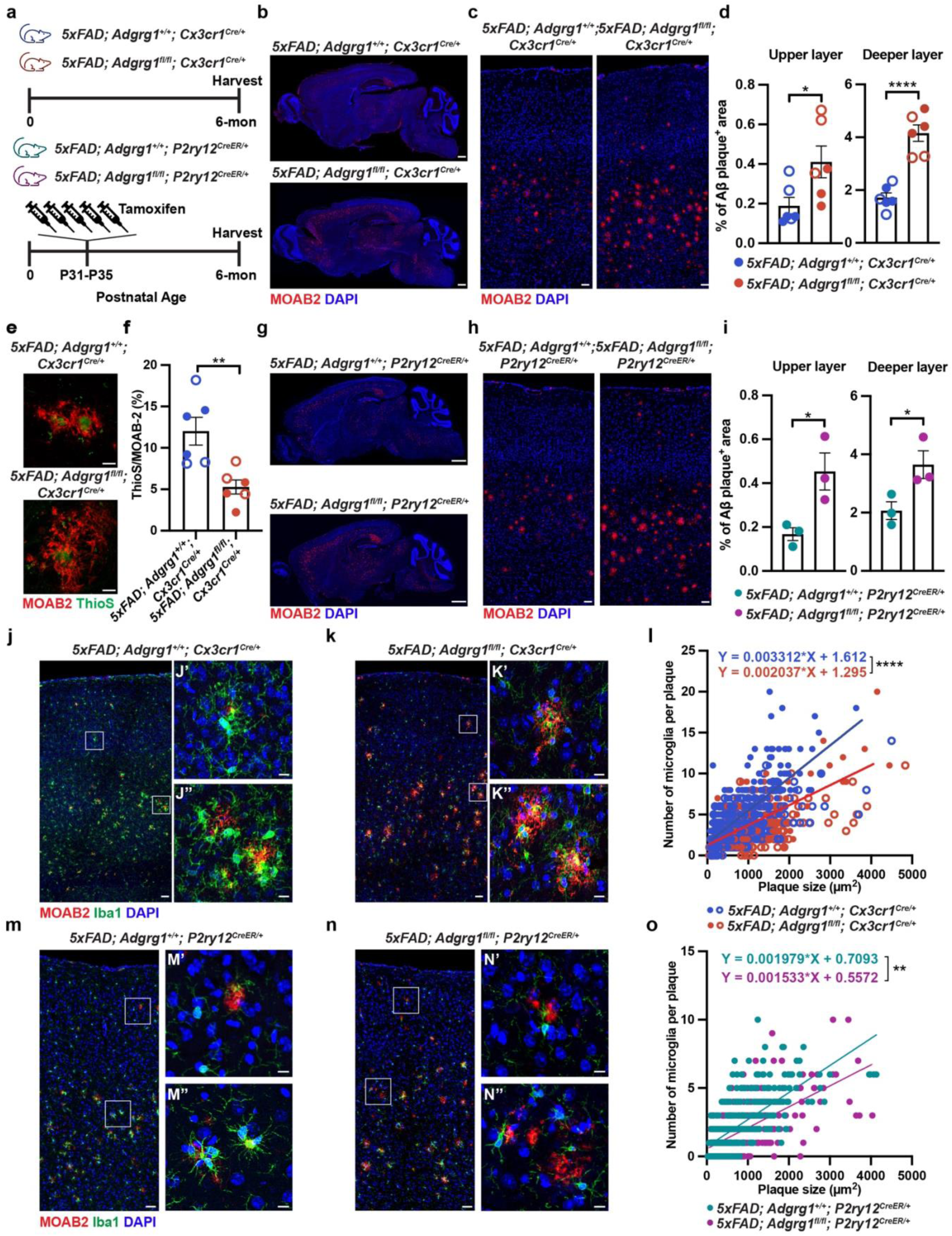
Conditional deletion of *Adgrg1* in microglia exacerbated amyloid deposition and diminished microglial capacity to respond to Aβ plaques. (a) Schematic of the experimental designs on constitutive *Cx3cr1^Cre/+^* mice and tamoxifen-inducible *P2ry12^CreER/+^* mice. (b) Representative tile scan images of Aβ staining (MOAB2) in 6-month-old *5xFAD; Adgrg1^+/+^; Cx3cr1^Cre/+^* and *5xFAD; Adgrg1^fl/fl^; Cx3cr1^Cre/+^* mice. Scale bar, 1 mm. (c) Representative images of MOAB2 staining in the cortices of 6-month-old *5xFAD; Adgrg1^+/+^; Cx3cr1^Cre/+^* and *5xFAD; Adgrg1^fl/fl^; Cx3cr1^Cre/+^* mice. Scale bar, 50 μm. (d) Percentage of MOAB2 positive area in the upper and deeper cortical layer of each genotype at 6months. Each circle represents one animal. * p=0.0349, **** p<0.0001. Unpaired two-tailed t-test. (e) Representative images of MOAB2 (red) and compact plaque core labeled by Thioflavin S (ThioS, green) in the cortices of 6-month-old mice. Scale bar, 10 μm. (f) Percentage of ThioS^+^ compact core area in the total plaque area. Each circle represents one animal. ** p=0.0050. Unpaired two-tailed t-test. (g) Representative tile scan images of Aβ staining (MOAB2, red) in 6-month-old *5xFAD; Adgrg1^+/+^; P2ry12^CreER/+^* and *5xFAD; Adgrg1^fl/fl^; P2ry12^CreER/+^* mice. Scale bar, 1 mm. (h) Representative immunofluorescent images of cortices stained for MOAB2 (red) and DAPI. Scale bar, 50 μm. (i) Percentage of MOAB2 positive area in the upper and deeper cortical layer of each genotype at 6-month-old. Each circle represents one animal. * p=0.0332 (for upper layer) * p=0.0489 (for deeper layer). Unpaired two-tailed t-test. (j-k) Representative images of cortices from 6-month-old *5xFAD; Adgrg1^+/+^; Cx3cr1^Cre/+^* and *5xFAD; Adgrg1^fl/fl^; Cx3cr1^Cre/+^*mice stained for MOAB2 (red), Iba1 (green) and DAPI. Zoom-in images indicating the upper cortical layers (j’ and k’) and deeper cortical layers (j’’ and k’’). Scale bar, 50 μm. (l) Quantification of microglia density within a 30-μm radius from the plaque cores and its relationship to the plaque size. Each circle represents single plaque analysed. 567 plaques for *5xFAD; Adgrg1^+/+^; Cx3cr1^Cre/+^*, 471 plaques for *5xFAD; Adgrg1^fl/fl^; Cx3cr1^Cre/+^*, from n=6 animals per genotype. **** p<0.0001, simple linear regression of slopes. (m-n) Representative images of cortices from 6-month-old *5xFAD; Adgrg1^+/+^; P2ry12^CreER/+^* and *5xFAD; Adgrg1^fl/fl^; P2ry12^CreER/+^*mice with tamoxifen induction at 1 month and stained for MOAB2 (red), Iba1 (green), and DAPI. Zoom-in images indicating the upper cortical layers (m’ and n’) and deeper cortical layers (m’’ and n’’). Scale bar, 50 μm. (o) Quantification of microglia density within a 30-μm radius from the plaque cores and its relationship to the plaque size. Each circle represents single plaque analysed. 379 plaques for *5xFAD; Adgrg1^+/+^; P2ry12^CreER/+^*, 430 plaques for *5xFAD; Adgrg1^fl/fl^; P2ry12^CreER/+^*, from n=3 animals per genotype. ** p=0.0012, simple linear regression of slopes. Filled circles, male mice; open circles, female mice. Data are presented as mean ± SEM. See also Figure S1.

We have previously shown that *Adgrg1^fl/fl^;Cx3cr1^Cre/+^*mice exhibit synaptic pruning and interneuron development deficits. ^13,14^ To alleviate any potential developmental impact of microglial *Adgrg1* deletion, we created an inducible microglial-specific knockout line by employing *P2ry12^CreER/+^;Ai14* mice^21^ to generate *5xFAD;Adgrg1^fl/fl^; P2ry12^CreER/+^;Ai14* (AD-icKO) and the *5xFAD;Adgrg1^+/+^;P2ry12^CreER/+^;Ai14* mice (AD-iCon) mice. Tamoxifen was administered at P31-P35 to induce *Adgrg1* deletion prior to amyloid plaque formation (**Figure 1a**).^17^ Five consecutive intraperitoneal tamoxifen injections (100 mg/kg) resulted in ∼90% recombination efficiency (**Figures S1f-j**). Our recombination rate exceeded what was observed in published reports, likely due to a favorable inter-loxP distance.^22,23^ Importantly, we observed increased Aβ load (**Figures 1g-i**, **S1k-l**) in 5xFAD-icKO mice compared to age-matched 5xFAD-iCon mice, recapitulating the phenotype observed in constitutive knockout models.

To gain insights into how ADGRG1 modulates microglial response in Aβ pathology, we next evaluated the density of microglia surrounding amyloid plaques. We found that the number of microglia within a 30-μm radius from plaques was significantly reduced in both 5xFAD-cKO and 5xFAD-icKO mice at 6 months, compared to 5xFAD-Con and 5xFAD-iCon mice (**Figures 1j-o**). Taken together, these findings suggest that ADGRG1 is necessary for effective microglial responses to limit amyloid deposition.

### Deleting microglial *Adgrg1* results in increased neuronal and synaptic loss and impaired neuritic integrity in 5xFAD mice

It has been reported that fibrillar Aβ_1-42_ disrupts synaptic function and contributes to neurotoxicity.^24^ To assess neuronal density across cortical layers, we performed NeuN^25^ immunostaining and observed a significant reduction of layer V neurons in 5xFAD-cKO mice compared to 5xFAD-Con (**Figures 2a-b**). To further analyze deeper layer cortical neurons, we conducted double-labeling of CTIP2^26^ and TBR1^27^, and found a significant reduction in CTIP2 density in layer V in the absence of microglial *Adgrg1* (**Figures 2c-d**). This phenotype was consistent between constitutive (**Figures 2a-d**) and inducible (**Figures 2e-h**) microglial *Adgrg1* deletion mouse models.

**Figure 2.**
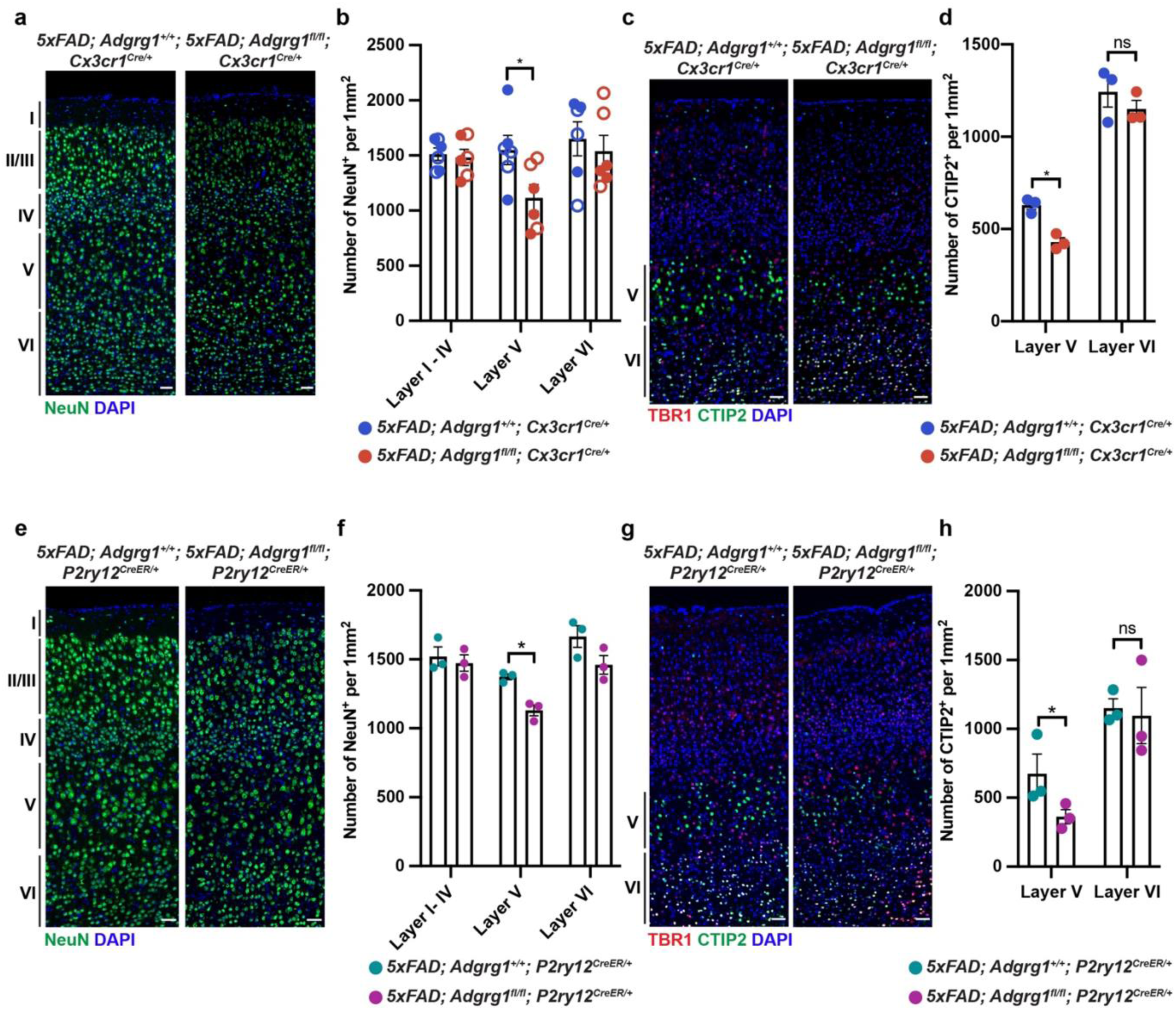
Microglial *Adgrg1* deficiency increased neuronal loss in 5xFAD mice. (a) Representative images of NeuN (green) staining in the cortices of 6-month-old *5xFAD; Adgrg1^+/+^; Cx3cr1^Cre/+^* and *5xFAD; Adgrg1^fl/fl^; Cx3cr1^Cre/+^* mice. Scale bar, 50 μm. (b) Quantification of NeuN-positive neurons in the cortices of 6-month-old mice. Each circle represents one animal. * p=0.0439. Two-way ANOVA with Bonferroni’s multiple comparisons test. (c) Representative images of the cortex with TBR1 (red) and CTIP2 (green) staining. Scale bar, 50 μm. (d) Quantification of the CTIP2-positive neurons in cortical layer V and layer VI in 6-month-old mice. Each circle represents one animal. * p=0.0459. Two-way ANOVA with Bonferroni’s multiple comparisons test. (e) Representative images of NeuN staining in the cortices of 6-month-old *5xFAD; Adgrg1^+/+^; P2ry12^CreER/+^* and *5xFAD; Adgrg1^fl/fl^; P2ry12^CreER/+^*mice. Scale bar, 50 μm. (f) Quantification of NeuN-positive neurons. Each circle represents one animal. * p=0.0384. Two-way ANOVA with Bonferroni’s multiple comparisons test. (g) Representative images of the cortex with TBR1 (red) and CTIP2 (green) staining. Scale bar, 50 μm. (h) Quantification of the CTIP2-positive neurons in cortical layer V and layer VI. Each circle represents one animal. * p=0.0479. Two-way ANOVA with Bonferroni’s multiple comparisons test. Filled circles, male mice; open circles, female mice. Data are presented as mean ± SEM.

Diffuse Aβ plaques are associated with increased neuritic dystrophy.^28^ To investigate the impact of microglial *Adgrg1* deletion on neuritic integrity, we performed MAP2^29,30^ staining in 6-month-old mouse brains and observed significantly shorter dendrites when *Adgrg1* was deleted (**Figures 3a-d**). LAMP1 staining, which marks amyloid plaque-associated axonal spheroids,^31–33^ showed increased immunoreactivity in both 5xFAD-cKO and 5xFAD-icKO mice relative to their respective controls (**Figures 3e-h**). At the synaptic level, co-labeling with the pre-synaptic marker vGlut2 and the post-synaptic marker PSD95 revealed a significant reduction in synaptic density in the cortical peri-plaque regions of 5xFAD-cKO mice and 5xFAD-icKO mice compared to their respective controls (**Figures 3i-l**). Taken together, these findings suggest that microglial ADGRG1 is crucial for maintaining neuronal health and synaptic integrity in 5xFAD mice.

**Figure 3.**
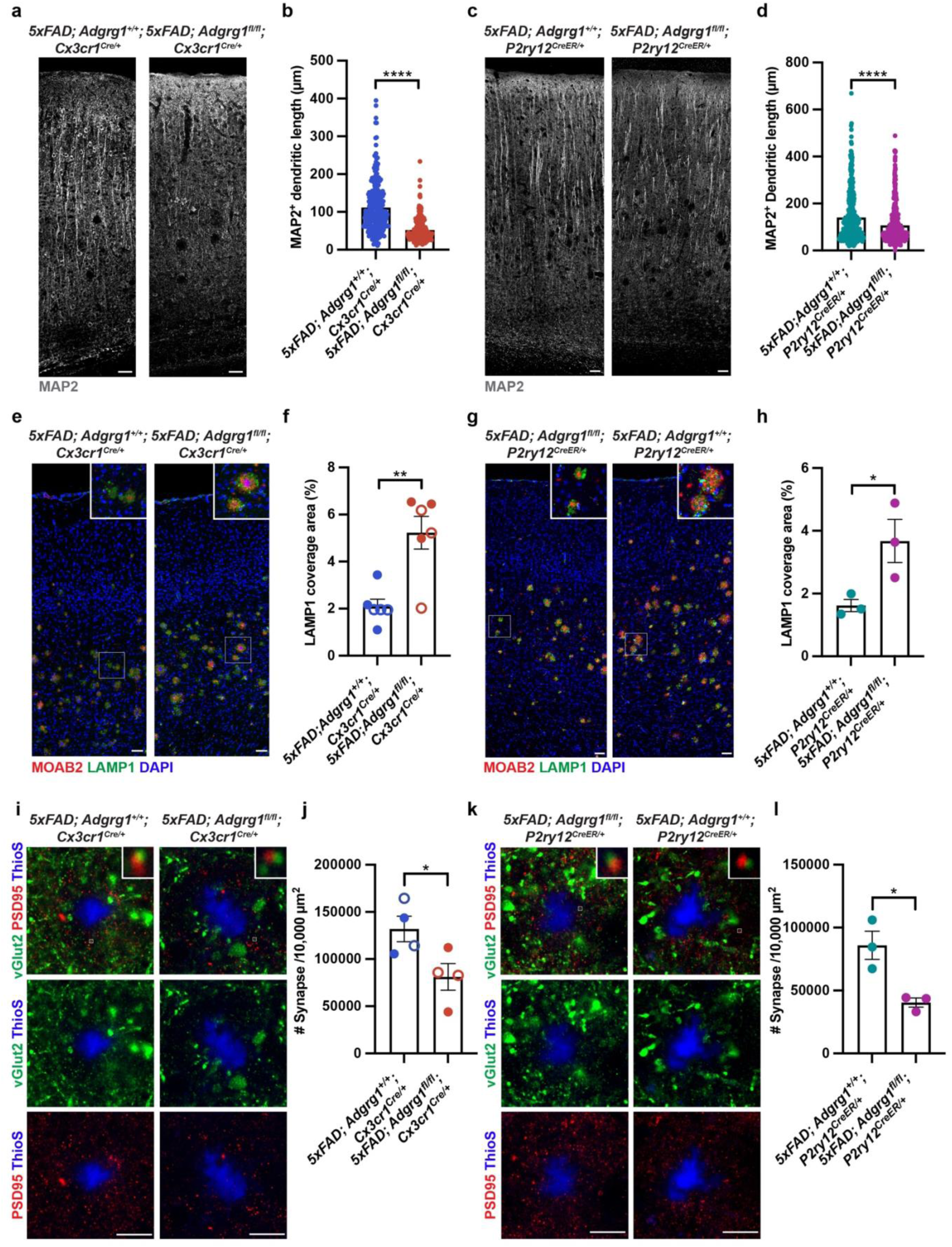
Loss of ADGRG1 in microglia impaired neuritic integrity and synaptic density. (a) Representative images of MAP2 staining in the cortices of 6-month-old *5xFAD; Adgrg1^+/+^; Cx3cr1^Cre/+^* and *5xFAD; Adgrg1^fl/fl^; Cx3cr1^Cre/+^* mice. Scale bar, 50 μm. (b) Quantification of MAP2^+^ dendritic length in the cortices of 6-month-old mice. Each circle represents one dendrite from n=6 animals per genotype. **** p<0.0001. Unpaired two-tailed t-test. (c) Representative images of MAP2 staining in the cortices of 6-month-old *5xFAD; Adgrg1^+/+^; P2ry12^CreER/+^* and *5xFAD; Adgrg1^fl/fl^; P2ry12^CreER/+^* mice. Scale bar, 50 μm. (d) Quantification of MAP2^+^ dendritic length in the cortices of 6-month-old mice. Each circle represents one dendrite from n=3 animals per genotype. **** p<0.0001. Unpaired two-tailed t-test. (e) Representative immunostaining images of cortex from 6-month-old mice stained for MOAB2 (red), LAMP1 (green), and DAPI. Scale bar, 50 μm. (f) Percentage of cortical LAMP1^+^ dystrophic neurites coverage area. Each circle represents one animal. * p=0.0020. Unpaired two-tailed t-test. Filled circles, male mice; open circles, female mice. (g) Representative images of MOAB2 and LAMP1 staining in 6-month-old *5xFAD; Adgrg1^+/+^; P2ry12^CreER/+^* and *5xFAD; Adgrg1^fl/fl^; P2ry12^CreER/+^* mice. Scale bar, 50 μm. (h) Percentage of cortical LAMP1^+^ coverage area. Each circle represents one animal. * p=0.0450. Unpaired two-tailed t-test. (i) Representative images of ThioS (blue), vGlut2 (green) labeling presynaptic terminals and PSD95 (red) labeling postsynaptic terminals in the cortices of 6-month-old *5xFAD; Adgrg1^+/+^; Cx3cr1^Cre/+^* and *5xFAD; Adgrg1^fl/fl^; Cx3cr1^Cre/+^* mice. Box indicating the zoom-in view of overlapped vGlut2 and PSD95 as synapses. Scale bar, 10 μm. (j) Quantification of synapse density. Each circle represents one animal. * p=0.0406. Unpaired two-tailed t-test. Filled circles, male mice; open circles, female mice. (k) Representative images of ThioS (blue), vGlut2 (green) and PSD95 (red) in the cortices of 6-month-old *5xFAD; Adgrg1^+/+^; P2ry12^CreER/+^* and *5xFAD; Adgrg1^fl/fl^; P2ry12^CreER/+^* mice. Box indicating the zoom-in view of overlapped vGlut2 and PSD95 as synapses. Scale bar, 10 μm. (l) Synapse density quantification. Each circle represents one animal. * p=0.0178. Unpaired two-tailed t-test. Data are presented as mean ± SEM.

### Deletion of *Adgrg1* in microglia accelerates memory deficits in 5xFAD mice

5xFAD mice manifest cognitive deficits at 6 months of age.^17,34,35^ Based on the worsened pathology associated with microglial *Adgrg1* deletion (**Figures 1-3**) and the similar phenotypes between the constitutive and tamoxifen-inducible knockout lines, we performed a series of behavioral tests in 4-month-old Con, *Adgrg1^fl/fl^;Cx3cr1^Cre/+^* (cKO), 5xFAD-Con, and 5xFAD-cKO mice (**Figure 4a**). We observed reduced body weight in male 5xFAD-cKO mice compared to Con mice (**Figures S2a-b**). Motor learning and function, as assessed by the rotarod test, were unaffected by *Adgrg1* deletion (**Figure 4b**). Consistent with the literature,^36^ 5xFAD mice exhibited longer latencies to fall from the rotarod than wild-type mice (**Figure 4b**).

**Figure 4.**
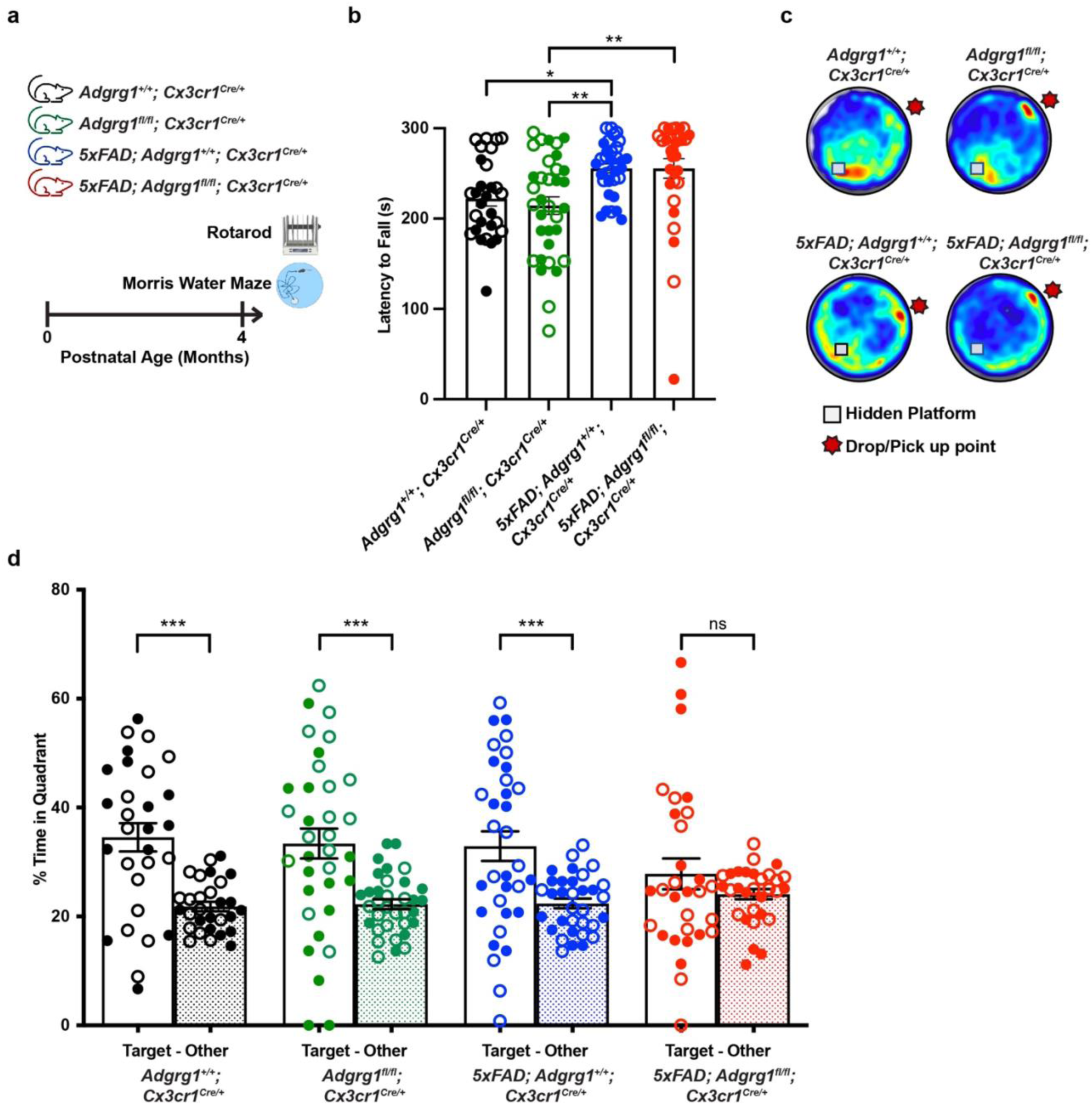
ADGRG1 deficiency in microglia led to learning and memory deficits. (a) Schematic of behavioral experimental design on 4-month-old mice. Schematic created with BioRender. (b) Rotarod test on 4-month-old mice from each genotype, shown as the maximum time that mice remained on the accelerating rotating rod. * p=0.0499, ** p=0.0050 (*Adgrg1^fl/fl^; Cx3cr1^Cre/+^* vs. *5xFAD; Adgrg1^+/+^; Cx3cr1^Cre/+^*), ** p=0.0061 (*Adgrg1^fl/fl^; Cx3cr1^Cre/+^*vs. *5xFAD; Adgrg1^fl/fl^; Cx3cr1^Cre/+^*). One-way ANOVA with Bonferroni’s multiple comparisons test. (c) 24-hour probe of Morris water maze test on 4-month-old mice from each genotype. Heat map images represent weighted occupancy across the entire 60-second trial. Hot colors indicate longer dwell times. The platform area is marked with a square. The star represents the drop/pick up point. (d) Percentage of time in target quadrant without submerged platform or in other quadrants. *** p=0.0001 (*Adgrg1^+/+^; Cx3cr1^Cre/+^*), *** p=0.0003 (*Adgrg1^fl/fl^; Cx3cr1^Cre/+^*), *** p=0.0009 (*5xFAD; Adgrg1^+/+^; Cx3cr1^Cre/+^*). Two-way ANOVA with Bonferroni’s multiple comparisons test. Sample size of behavioral tests: 4-month-old *Adgrg1^+/+^; Cx3cr1^Cre/+^* (n=29), *Adgrg1^fl/fl^; Cx3cr1^Cre/+^* (n=34), *5xFAD; Adgrg1^+/+^; Cx3cr1^Cre/+^* (n=33), and *5xFAD; Adgrg1^fl/fl^; Cx3cr1^Cre/+^*(n=31). Filled circles, male mice; open circles, female mice. Data are presented as mean ± SEM. See also Figure S2.

Spatial learning and memory were evaluated using the Morris water maze. During the memory recall session without a submerged platform, 4-month-old 5xFAD-cKO mice spent approximately equal amounts of time in the target and non-target quadrants, indicating impaired memory retention. In contrast, mice from other genotypes spent significantly more time in the target quadrant, demonstrating intact spatial memory (**Figures 4c-d**). Furthermore, 5xFAD-cKO mice exhibited a notably higher swimming speed compared to cKO mice, suggesting motor hyperactivity (**Figure S2c**). However, the average distance from the platform remained consistent across different genotypes (**Figure S2d**). Taken together, these behavioral tests support that, while sparing body weight and motor skills, microglial *Adgrg1* deletion significantly compromises spatial learning and memory capabilities in 5xFAD mice.

### *Adgrg1* deletion alters microglial response to Aβ at the transcriptomic level

To delineate the molecular mechanism by which ADGRG1 regulates microglial responses to amyloid deposition, we conducted snRNA-seq on nuclei isolated from the neocortex of 6-month-old Con, cKO, 5xFAD-Con, and 5xFAD-cKO mice (**Figures 5a and S3a-e**). Given that microglia are often under-represented in snRNA-seq datasets,^37^ we employed an enrichment strategy targeting glial cells. Nuclei were stained with DAPI and NeuN, followed by flow cytometry sorting for DAPI^+^ and DAPI^+^/NeuN^-^ populations (**Figure 5a**). Single-nucleus cDNA libraries were generated using the Chromium Next GEM v3.1 from 10X Genomics. A total of 191,292 single nuclei were sequenced, with a target of 50,000 reads per cell, resulting in the detection of 1500-2500 genes per cell on average. Eleven clusters were identified based on the transcriptomic profile as shown in the uniform manifold approximation projection (UMAP) plot (**Figure 5b**). The clusters were annotated based on marker genes identified of previous known cell type (**Figure S3e**). Microglial population were identified by shared enrichment of *Csf1r*, *Spi1*, *P2ry12*, and *Cx3cr1* (**Figure S3g**).

**Figure 5.**
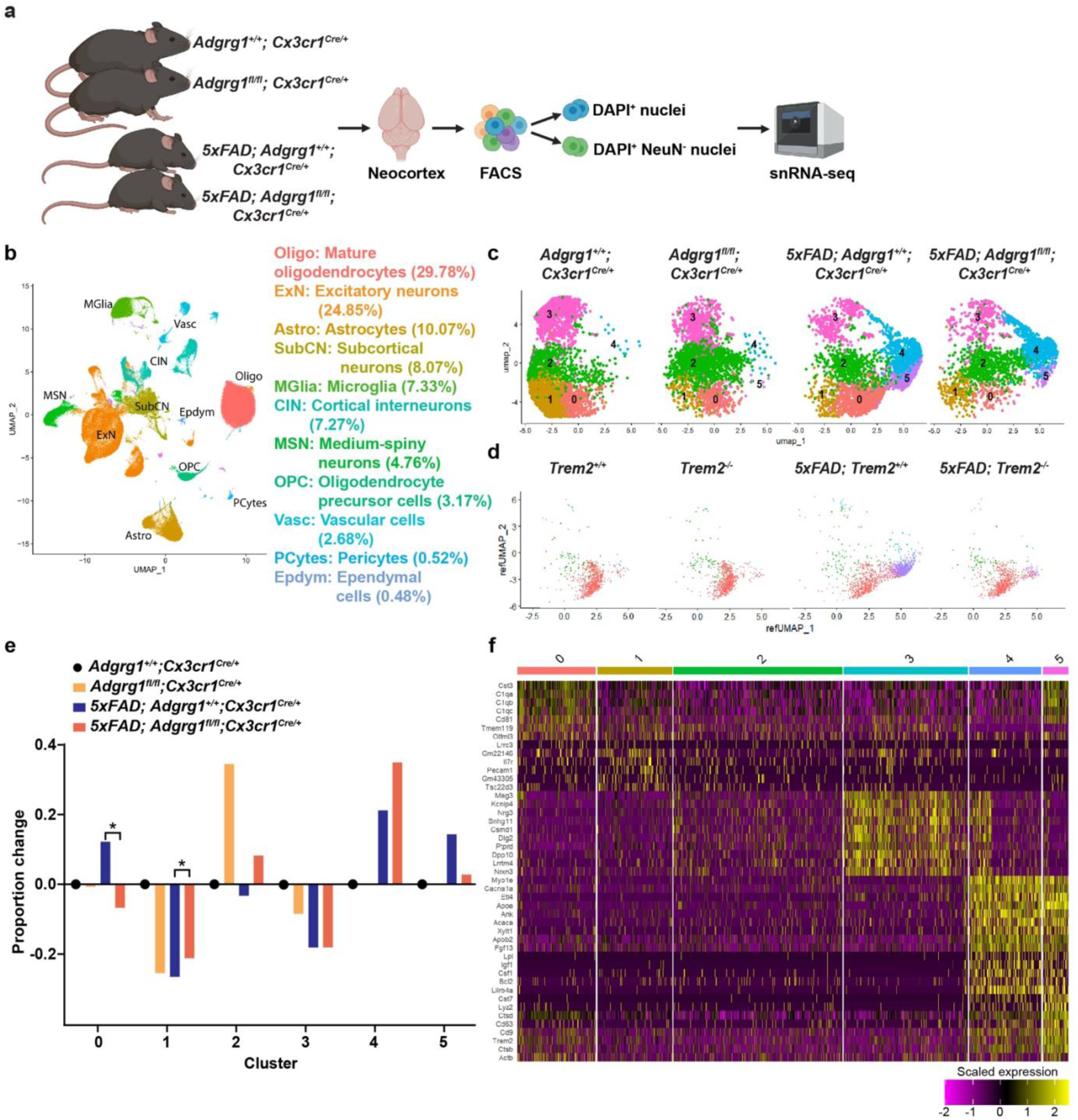
The deletion of *Adgrg1* in microglia altered microglial transcriptomic profiles. (a) Schematic of the snRNA-seq experimental design on 6-month-old mice (n=3 male and 3 female mice per genotype). Schematic created with BioRender.com. (b) UMAP plots of all single nuclei and their annotated cell types and proportion. (c) UMAP plots of microglia subclusters split by genotype and colored according to subclusters. (d) UMAP plots of mouse microglia snRNA-seq datasets from Zhou *et al.* (3,712 nuclei) re-analyzed and projected onto our microglial subclustering as in (c). (e) Proportion of microglia subclusters from each genotype, normalized to *Adgrg1^+/+^; Cx3cr1^Cre/+^.* * p=0.1418 for cluster 0, * p=0.1952 for cluster 1. Unpaired two-tailed t-test. (f) Gene expression heatmap showing the top enriched genes for each microglia cluster. See also Figure S3.

To further dissect the microglial population, we performed iterative clustering on 11,398 microglia cells and identified five transcriptomic subclusters. These subclusters were labeled according to the identified marker genes: homeostatic microglia (*Tmem119*, *P2ry12*; clusters 0-2), neuronal supportive microglia (*Nrxn3*^38^, *Cadm2*^39^; cluster 3), and disease-associated microglia (DAM; clusters 4, 5) (**Figures 5c-f**). Notably, DAMs were consistently found across 5xFAD-Con and 5xFAD-cKO mice (**Figure 5c**), indicating that *Adgrg1* deletion did not impair microglial progression to DAM states, unlike *Trem2* deletion in 5xFAD mice^40^ (**Figure 5d**). However, microglial *Adgrg1* deletion resulted in a reduction of clusters 0 and 5 and an increase in cluster 4 cells (**Figures 5c**, **5e and S3h**), leading to elevated expression of lipid processing and cell survival genes, including *Lpl*, *Igf1* and *Csf1* (cluster 4), and decreased expression of lysosomal genes such as *Lyz2*, *Ctsd*, *Tyrobp*, *Cst7* (cluster 5) (**Figures S3j-k**). Furthermore, we found that homeostatic microglia (*P2ry12*, *P2ry13*, *Cx3cr1*, *Tmem119*) with enhanced phagocytic function (*Cd81*, *C1qa/b/c*) were less abundant in 5xFAD-cKO compared to 5xFAD-Con mice (cluster 0, **Figures 5c**, **5e, 5f and S3i**). Taken together, these results highlight that microglial *Adgrg1* deletion leads to reduced levels of genes related to homeostasis, phagocytosis and lysosomal function, suggesting a loss of protective microglial responses.

### Deletion of *Adgrg1* diminishes microglial phagocytosis of Aβ *in vivo* and in human hESC-derived microglia

To further investigate how ADGRG1 influences microglial response to Aβ pathology, we performed differential gene expression (DEG) analysis and revealed a downregulation of homeostatic genes, such as *Tmem119*, *Cx3cr1,* and *P2ry12*, as well as genes associated with lysosomal function, such as *Cd68, Grn*, and several Cathepsin family members (*Ctsb*, *Ctsd*, *Ctss*, and *Ctsl*), in 5xFAD-cKO microglia (**Figure 6a**). In contrast, genes related to AD pathology, including Ankyrin (*Ank*) and Amyloid Beta Precursor Protein Binding Family B Member 2 (*Apbb2*), were found to be upregulated (**Figure 6a**). Gene ontology (GO) pathway analysis further highlighted that lysosome and phagocytosis pathways in microglia were significantly affected by *Adgrg1* deficiency (**Figure 6b**).

**Figure 6.**
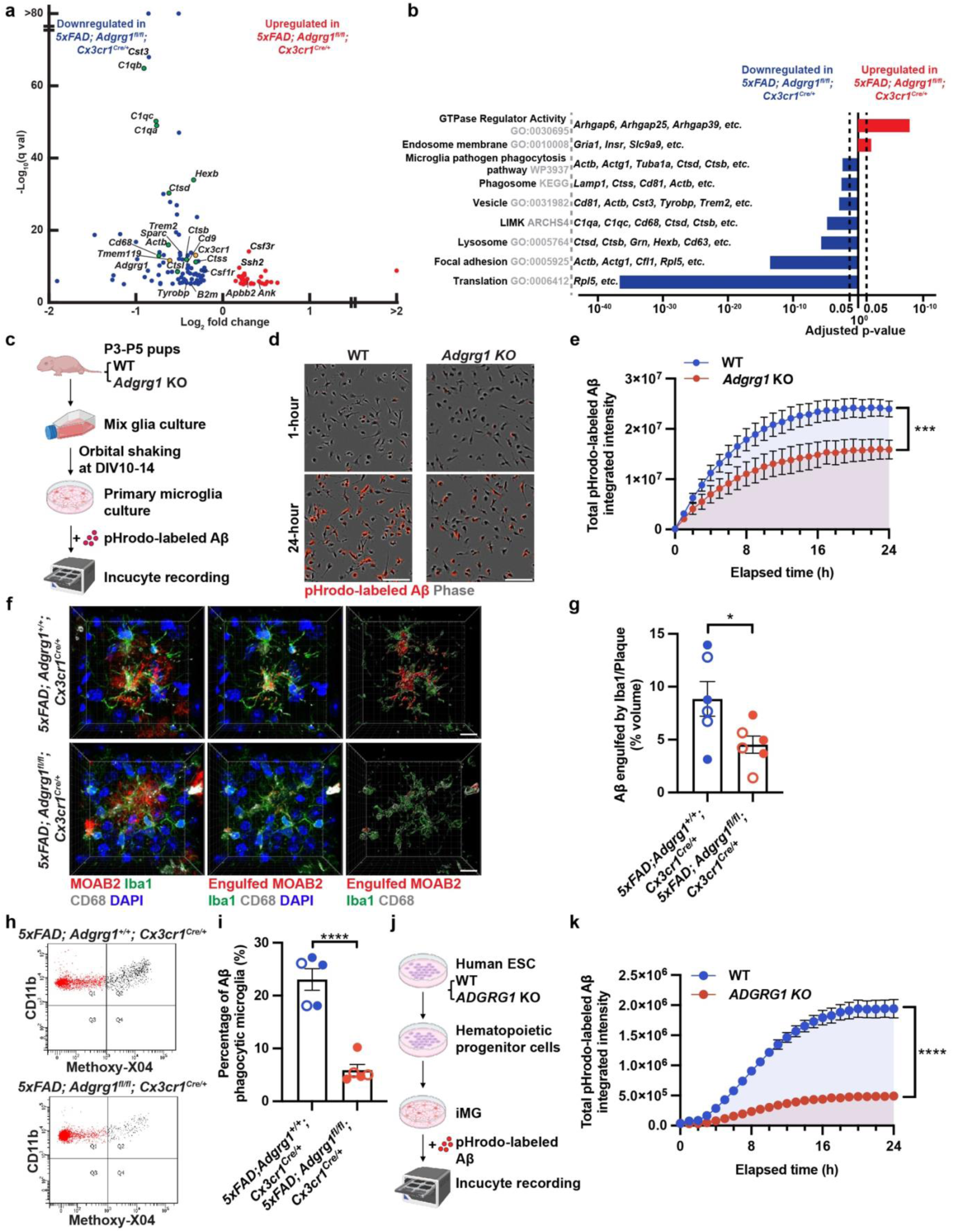
Microglial ADGRG1 regulates Aβ phagocytosis in AD mouse models and hESC-derived microglia. (a) Volcano plot showing significant differential expressed genes in microglia of *5xFAD; Adgrg1^+/+^; Cx3cr1^Cre/+^* versus *5xFAD; Adgrg1^fl/fl^; Cx3cr1^Cre/+^*. Genes related with microglial homeostasis are highlighted in yellow and phagocytosis and lysosomal pathways are in green. (b) Gene ontology analysis showing hallmark pathways based on the top 500 upregulated and downregulated DEGs in microglia. (c) Schematic showing primary microglia derived from wild-type (WT) and *Adgrg1* KO pups treated with pHrodo-labeled Aβ and recorded using the Incucyte S3 live imaging system. (d) Representative Incucyte images of WT and KO primary microglia phagocytose pHrodo-labeled Aβ after 1-hour and 24-hour incubation. Scale bar, 100 μm. (e) pHrodo-labeled Aβ signal (total integrated intensity) measured every hour in a time course of 24 hours. Quantification of area under curve. n=3 biological replicates per genotype. *** p=0.0008. Unpaired two-tailed t-test. (f) 3D reconstruction of microglia from 6-month-old *5xFAD; Adgrg1^+/+^; Cx3cr1^Cre/+^* and *5xFAD; Adgrg1^fl/fl^; Cx3cr1^Cre/+^* mice, showing engulfed Aβ inside Iba1^+^ microglia. Scale bar, 10 μm. (g) Percentage of the engulfed Aβ volume to the total Aβ volume. Each circle represents one animal. * p=0.0400. Unpaired two-tailed t-test. Filled circles, male mice; open circles, female mice. (h) Representative FACS plots of methoxy-X04^+^ plaque and CD11b^+^ microglia in the cortex of 6-month-old *5xFAD; Adgrg1^+/+^; Cx3cr1^Cre/+^* and *5xFAD; Adgrg1^fl/fl^; Cx3cr1^Cre/+^* mice. Q2 represents the Aβ plaques that were engulfed by microglia within the past three hours. (i) Percentage of methoxy-X04^+^ microglia. Each circle represents one animal. **** p<0.0001. Unpaired two-tailed t-test. Filled circles, male mice; open circles, female mice. (j) Schematic showing the differentiation, maturation, treatment and Incucyte recording of human WT and *ADGRG1* KO hESC-derived microglia. (k) pHrodo-labeled Aβ signal (total integrated intensity) in WT and *ADGRG1* KO hESC-derived microglia measured every hour in a time course of 24 hours. Quantification of area under curve. n=8 for WT, n=9 for *ADGRG1* KO. n represents the number of independent microglial treatments. **** p<0.0001. Unpaired two-tailed t-test. Data are presented as mean ± SEM. Schematic created with BioRender.com. See also Figure S4.

To connect these transcriptomic findings to functional outcomes, we asked whether ADGRG1 deficiency affected microglial phagocytosis. We first assessed pHrodo-labeled oligomeric Aβ_1-42_ uptake in primary cultured microglia (**Figure 6c**). We confirmed over 98% purity of our isolated primary microglia (**Figure S4a**) and the absence of ADGRG1 protein in *Adgrg1* KO microglia (**Figure S4b**). We employed Incucyte live-cell imaging system to record RFP signals hourly over a 24-hour duration in wild-type (WT) and *Adgrg1* KO microglia. *Adgrg1*-deficient microglia phagocytosed significantly less pHrodo-Aβ_1-42_ than WT microglia (**Figures 6d-e**). We next performed 3D reconstructions of Iba1, CD68 and MOAB2 staining on 6-month-old 5xFAD-Con and 5xFAD-cKO brains. 5xFAD-cKO microglia engulfed significantly less Aβ compared to the 5xFAD-Con microglia (**Figures 6f-g**). To quantify Aβ uptake *in vivo*, we injected 6-month-old mice intraperitoneally (i.p.) with Methoxy-X04, a fluorescent dye that specifically binds to Aβ fibrillar β-sheet structures,^41,42^ and harvested mouse brain cortices 3 hours later. Microglia were labeled with CD11b antibody for subsequent flow cytometry analysis (**Figures S4c-d**). 5xFAD-cKO mice had significantly fewer Methoxy-X04^+^ microglia, suggesting reduced Aβ engulfment with the deletion of microglial *Adgrg1* (**Figures 6h-i**).

To investigate whether the phagocytosis impairment observed in *Adgrg1*-deficient mouse microglia also occurs in human microglia, we utilized CRISPR-Cas9 to generate an *ADGRG1* KO hESC line (**Figure S4e**) and induced KO and isogenic control hESCs into microglia (iMG) (**Figures 6j and S4f**). These cells were treated with pHrodo-oligomeric Aβ. Microglial phagocytic activity was monitored hourly for a time course of 24 hours by Incucyte. Notably, *ADGRG1* KO iMG showed significantly reduced Aβ engulfment compared to WT iMG, indicating that ADGRG1 is required for human microglial phagocytosis (**Figure 6k**). Together, these findings demonstrate that ADGRG1 deficiency significantly impairs microglial phagocytosis of Aβ in both mouse and human microglia.

### ADGRG1 is highly expressed in individuals with mild cognitive impairment and positively correlates with phagocytosis-related genes in human microglia

Thus far, we have demonstrated that microglial ADGRG1 is critical for limiting Aβ plaque burden. To correlate our findings in mouse models with human AD, we first examined microglial ADGRG1 protein levels in human brain tissues (**Table S1**). We performed immunostaining for Iba1, CG4 (a monoclonal antibody against human ADGRG1),^43^ and the Aβ antibody H31L21^44^ on human middle temporal gyrus tissue sections, a brain region vulnerable to AD pathological changes.^45–49^ We observed that microglia from individuals with mild cognitive impairment due to AD (MCI, AD Thal phase ≤4, Braak stage ≤4, AD neurologic change low to intermediate) exhibited prominent CG4-positive signals compared to those from AD patients, suggesting a potential resilient function of microglial ADGRG1 in individuals with MCI (**Figures 7a-b**).

**Figure 7.**
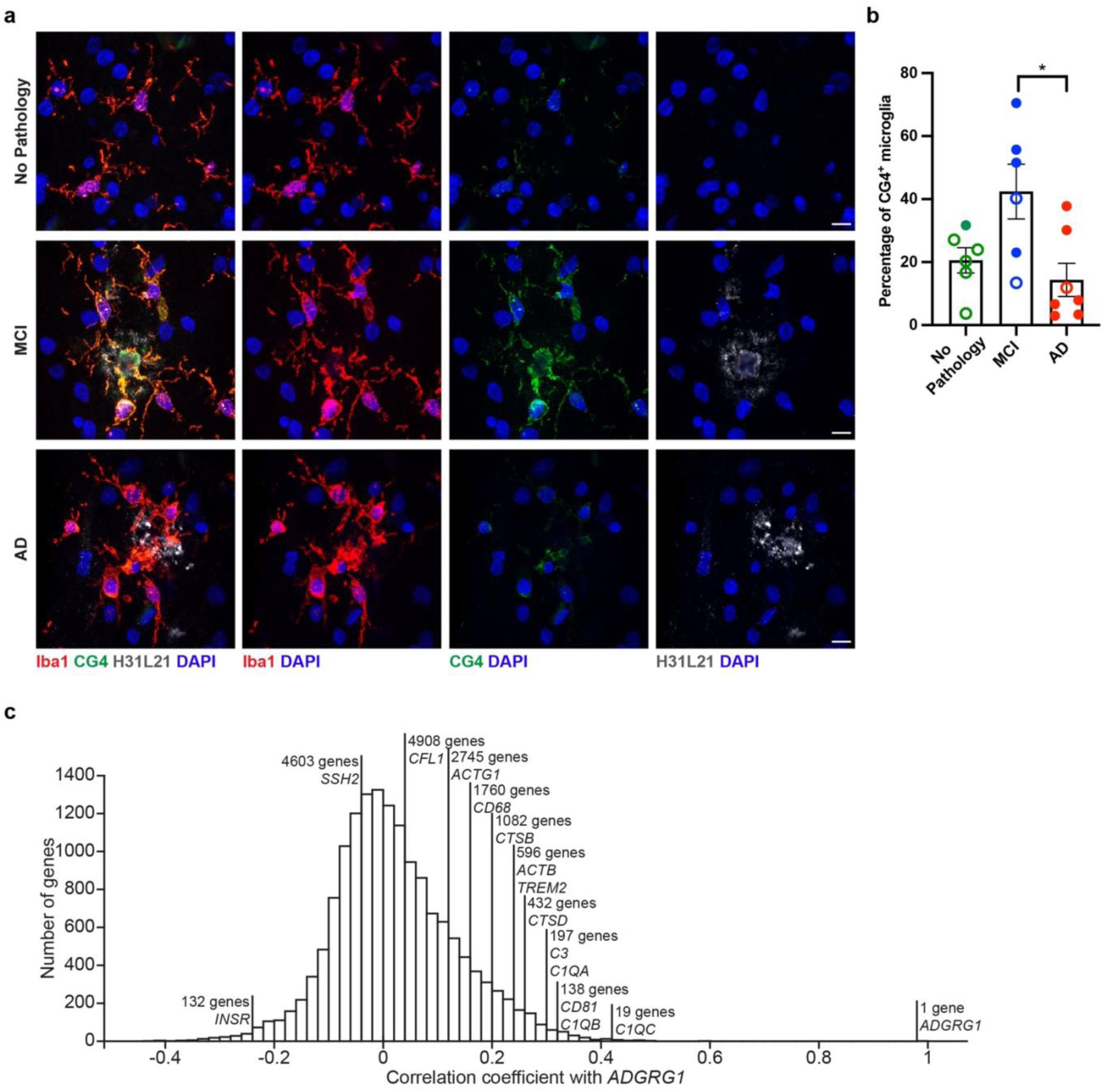
ADGRG1 expression and correlation with phagocytosis-related genes in human microglia. (a) Representative immunofluorescence images of human temporal gyrus sections stained for Iba1 (red), CG4 (ADGRG1, green), H31L21 (Aβ, gray), and DAPI (blue) in aged-matched no pathology, mild-cognitive impairment (MCI) and AD subjects. Scale bar, 10 μm. (b) Quantification of percentage of Iba1^+^ microglia positive for CG4. Each circle represents individual subject. * p=0.0159. One-way ANOVA with Bonferroni’s multiple comparison. Filled circles, male; open circles, female. (c) Histogram of Pearson correlation coefficient demonstrating the linear relationship between microglia *ADGRG1* expression and other microglial genes. Data are presented as mean ± SEM. See also Figure S5 and Table S1.

Given the unavailability of natural loss-of-function mutation data for *ADGRG1* in existing human AD samples, we adopted an alternative approach to assess the correlation of *ADGRG1* expression with other genes in human microglia of individuals with AD. We performed a transcriptomic meta-analysis across 590 human brain tissues using data from published snRNA-seq, scRNA-seq^50–53^ and the Religious Orders Study and Rush Memory and Aging Project (ROSMAP) studies.^54^ We found that microglial *ADGRG1* expression positively correlated with the expression of phagocytosis-related genes, such as *CD68*, *CTSD*, *CTSB*, and negatively correlated with *SSH2* and *INSR*, aligning with findings from our mouse models (**Figure 7c**). Additionally, an analysis of GWAS data on AD risk genes (GWASdb) revealed that *ADGRG1* is one of the top 300 significantly altered genes in AD.^55^ In a comparison of microglial DEGs from our snRNA-seq data with known human AD risk genes, 24 AD risk genes overlapped with DEGs upregulated in 5xFAD-Con microglia, whereas 97 risk genes overlapped with those upregulated in 5xFAD-cKO, suggesting a substantial increase in the overlap of DEGs with known AD risk genes due to *ADGRG1* deficiency (**Figure S5**). Taken together, these data demonstrate the linkage between microglial ADGRG1 with phagocytosis-related as well as AD risk genes in human, indicating ADGRG1’s potential role in modulating disease resilience via functional properties of microglia.

## Discussion

In this study, we demonstrate the crucial role of a G protein-coupled receptor ADGRG1 in regulating protective microglial responses to amyloid plaques in AD pathology. We find that ADGRG1 deficiency compromises the ability of microglia to associate with and phagocytose Aβ, leading to increased amyloid deposition in the cortex and hippocampus. Through various assays, including primary microglia cultures, immunostaining and subsequent 3D reconstructions, *in vivo* phagocytosis assays, and hESC-derived microglial phagocytosis assays, we demonstrate that ADGRG1 is required for microglial phagocytosis of Aβ. Microglia are highly heterogeneous immune cells in the brain and display remarkable plasticity in response to environmental cues during development, disease, and aging. They adapt their functions through distinct phenotypic states driven by unique, sometimes overlapping, transcriptomic profiles.^6,56^ During neurodegenerative disease conditions such as AD, microglia transition into DAM states by first downregulating homeostatic genes like *Cx3cr1*, *P2ry12*, and *Tmem119* in stage 1 DAM, and subsequently upregulating genes such as *Tyrobp*, *Apoe*, and *Trem2* in stage 2 DAM to enhance their phagocytic capacity under the TREM2-SYK regulation.^40,57–59^ Additionally, axon tract microglia (ATM)^60^ and proliferative region-associated microglia (PAM)^61^ are observed during development, playing roles in axonal maintenance, and supporting neurogenesis and oligodendrocyte differentiation, respectively. Our findings reveal that microglial *Adgrg1* deletion in 5xFAD mice leads to a unique transcriptomic profile featured by a downregulation of both homeostatic and phagocytic genes, indicating a dysfunctional microglial state that exacerbates Aβ deposition and neurodegeneration. Deletion of microglial *Adgrg1* leads to a significant reduction in the microglial subcluster cluster 0, marked by homeostatic microglia with enhanced phagocytic function. These findings advance our understanding of how microglial states are modulated through membrane GPCRs. Unlike *Trem2* deletions, where microglia fail to fully transition from homeostatic to DAM states,^40^ *Adgrg1* deletion shifts microglial function without affecting DAM progression but alters subpopulations within this state. Specifically, there is an increased expression of lipid processing genes and decreased expression of lysosomal genes, suggesting a potential link to lipid-droplets accumulating microglia (LDAM) observed in aging^62^ and AD,^63^ particularly among patients with the *APOE4/4* genotype.

Our published work and the present study uncover the dual functionality of microglial ADGRG1 deletion on synaptic density across different stages: early neurodevelopment and neurodegeneration. During development, inactive synapses externalize phosphatidylserine (PS) on the presynaptic terminal, facilitating microglial ADGRG1-mediated synaptic pruning. Deleting microglial *Adgrg1* during this stage results in an accumulation of excess synapses.^14^ Conversely, in 5xFAD mice, specifically around the peri-plaque regions, we observed a reduction in synaptic density following microglial *Adgrg1* knockout, likely attributable to the neurotoxic effects of Aβ plaques in 5xFAD mice. In both 5xFAD-cKO and 5xFAD-icKO brains, a higher amyloid burden is observed due to impaired microglial function, which results in worsened neuritic dystrophy. It is reported that Aβ-related hyperactive synapses externalize PS, resulting in synaptic loss in AD mouse models.^64–66^

In summary, our results reveal the previously unexplored role of microglial ADGRG1 in influencing microglial responses in AD, providing new insights into how a specific GPCR shapes microglial behavior and disease progression. Future research needs to explore whether enhancing ADGRG1 expression in microglia can serve as a therapeutic strategy to promote Aβ clearance and mitigate AD progression. By manipulating ADGRG1 expression or its downstream signaling pathways, one can potentially guide microglia towards a protective state in AD.

## Acknowledgements

We thank Neurodegenerative Disease Brain Bank at the University of California, San Francisco, which receives funding support from NIH grants P01AG019724 and P50AG023501, the Consortium for Frontotemporal Dementia Research, and the Tau Consortium. We are grateful to Piao Lab members for helpful comments. We thank Dr. Eric Huang for thoughtful comments on the manuscript, Dr. Yu-Hsin Huang for the support on primary microglia culture, Dr. Wendell Lim for access to the Incucyte, Dr. Thomas Arnold for his sharing of *P2ry12^CreER/+^;Ai14* mice. We acknowledge the following funding support: to X.P.: NIH/NINDS (R01NS094164 and R01NS108446) and Alzheimer’s Association (23AARG-NTF-1030341); to B.Z.: NIA (K99AG081694) and the BrightFocus foundation postdoctoral fellowship (A2021020F).

## Author contributions

Conceptualization by X.P. and B.Z.; *In vitro and in vivo* studies by B.Z., D.Y., T.L., R.S., S.F. and L.C.; Bioinformatic analysis by A.W. with help from T.N.; Experiments on human AD samples supervised by W.S., L.G. and S.P; hESC generation by Y.P. under the supervision of A.K., Manuscript writing: B.Z., R.R. and X.P.. All authors read and edited the manuscript.

## Declaration of interests

The authors declare no competing interests.

---

**Figure S1.**
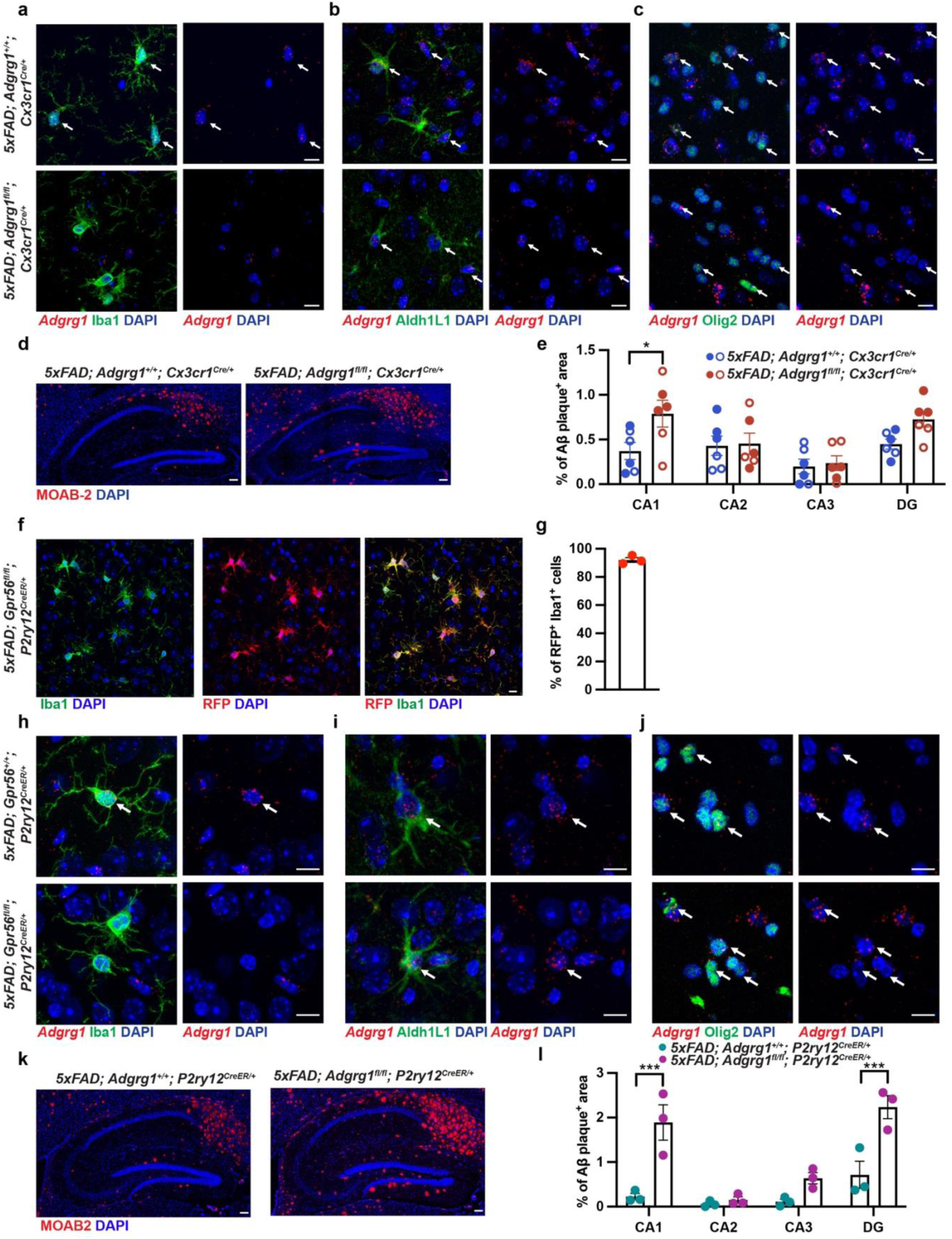
Microglial-specific deletion of *Adgrg1* and its consequences on amyloid deposition in hippocampus, related to Figure 1. (a-c) Representative immunofluorescence images of cortices from 6-month-old *5xFAD; Adgrg1^+/+^; Cx3cr1^Cre/+^* and *5xFAD; Adgrg1^fl/fl^; Cx3cr1^Cre/+^* mice stained for *Adgrg1* transcripts by RNAscope (red), different glial cell type marker (green), and DAPI. Arrows indicate *Adgrg1* transcripts-positive cells. Scale bar, 10 μm. (d) Representative images of amyloid staining (MOAB2, red) in the hippocampi of 6-month-old *5xFAD; Adgrg1^+/+^; Cx3cr1^Cre/+^*and *5xFAD; Adgrg1^fl/fl^; Cx3cr1^Cre/+^* mice. Scale bar, 100 μm. (e) Quantification of Aβ load in hippocampal CA1, CA2, CA3 and dentate gyrus (DG) subregions. Each circle represents one animal. * p=0.0225. Two-way ANOVA with Bonferroni’s multiple comparison. Filled circles, male; open circles, female. (f) Representative immunofluorescence labeling of Iba1 (green), RFP (red), and DAPI in the cortices of 6-month-old *5xFAD; Adgrg1^fl/fl^; P2ry12^CreER/+^* mice with tamoxifen induction at P31-P35. Scale bar, 10 μm. (g) Percentage of cells double-positive for RFP and Iba1, relative to the total number of Iba1-positive cells. Each circle represents one animal. (h-j) Representative images of cortices from 6-month-old *5xFAD; Adgrg1^+/+^; P2ry12^CreER/+^* and *5xFAD; Adgrg1^fl/fl^; P2ry12^CreER/+^* stained for *Adgrg1* transcripts by RNAscope (red), different glial cell type marker (green), and DAPI. Arrows indicate *Adgrg1* transcripts-positive cells. Scale bar, 10 μm. (k) Representative images of Aβ staining (MOAB2, red) in the hippocampi of 6-month-old *5xFAD; Adgrg1^+/+^; P2ry12^CreER/+^*and *5xFAD; Adgrg1^fl/fl^; P2ry12^CreER/+^* mice. Scale bar, 100 μm. (l) Quantification of Aβ load in hippocampal CA1, CA2, CA3 and DG subregions. Each circle represents one animal. *** p=0.0002 (CA1), *** p=0.0004 (DG). Two-way ANOVA with Bonferroni’s multiple comparison. Data are presented as mean ± SEM.

**Figure S2.**
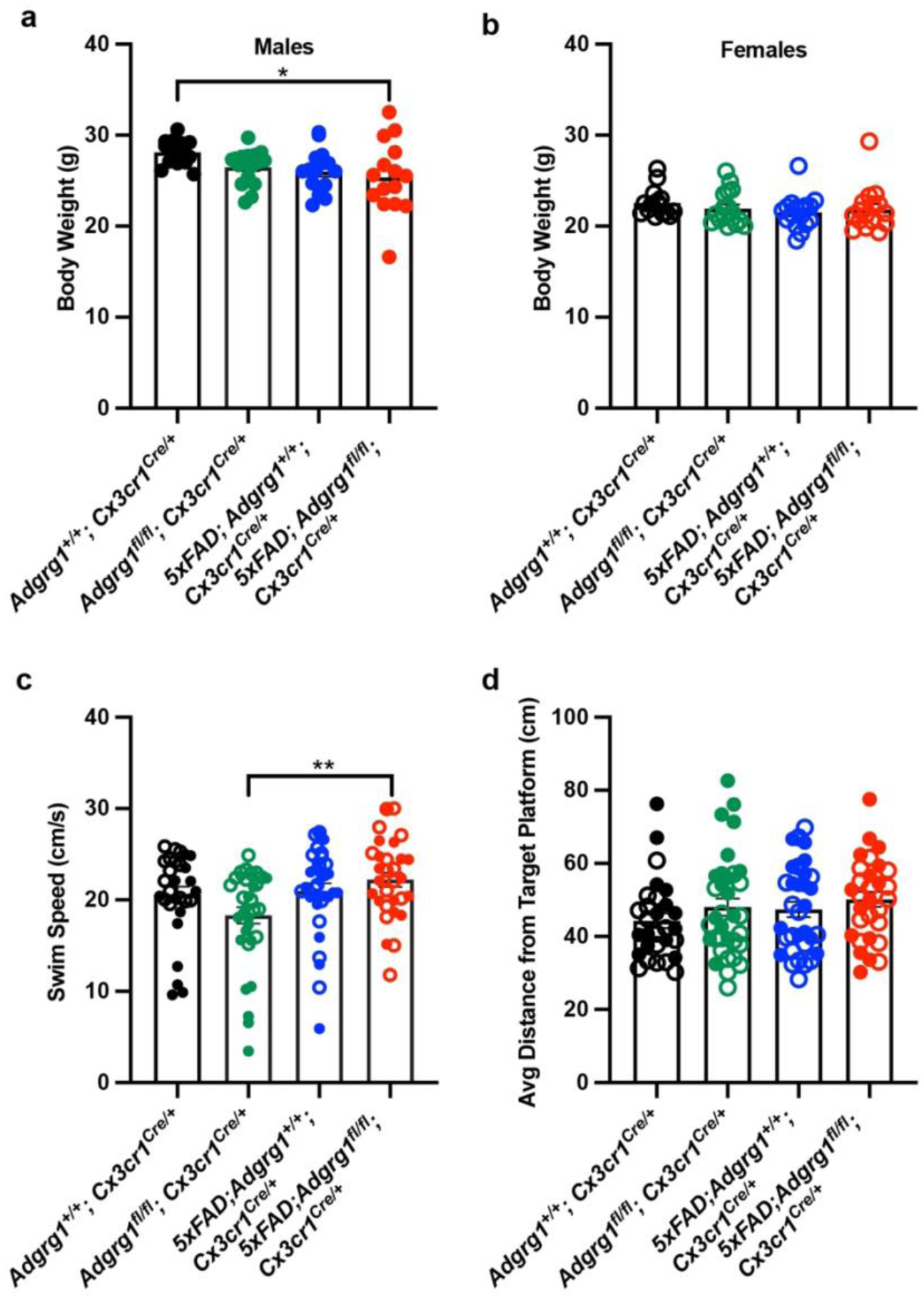
Body weight, swim speed and target proximity in Morris water maze, related to Figure 4. (a-b) Body weights of 4-month-old male and female mice at the beginning of the behavioral tests. Each circle represents one animal. * p=0.0310. One-way ANOVA with Bonferroni’s multiple comparison. (c) The average swimming speed during the Morris water maze 24-hour probe testing. Each circle represents one animal. ** p=0.0092. One-way ANOVA with Bonferroni’s multiple comparison. (d) The average distance from target platform during 24-hour probe testing. Each circle represents one animal. Filled circles, male mice; open circles, female mice. Data are presented as mean ± SEM.

**Figure S3.**
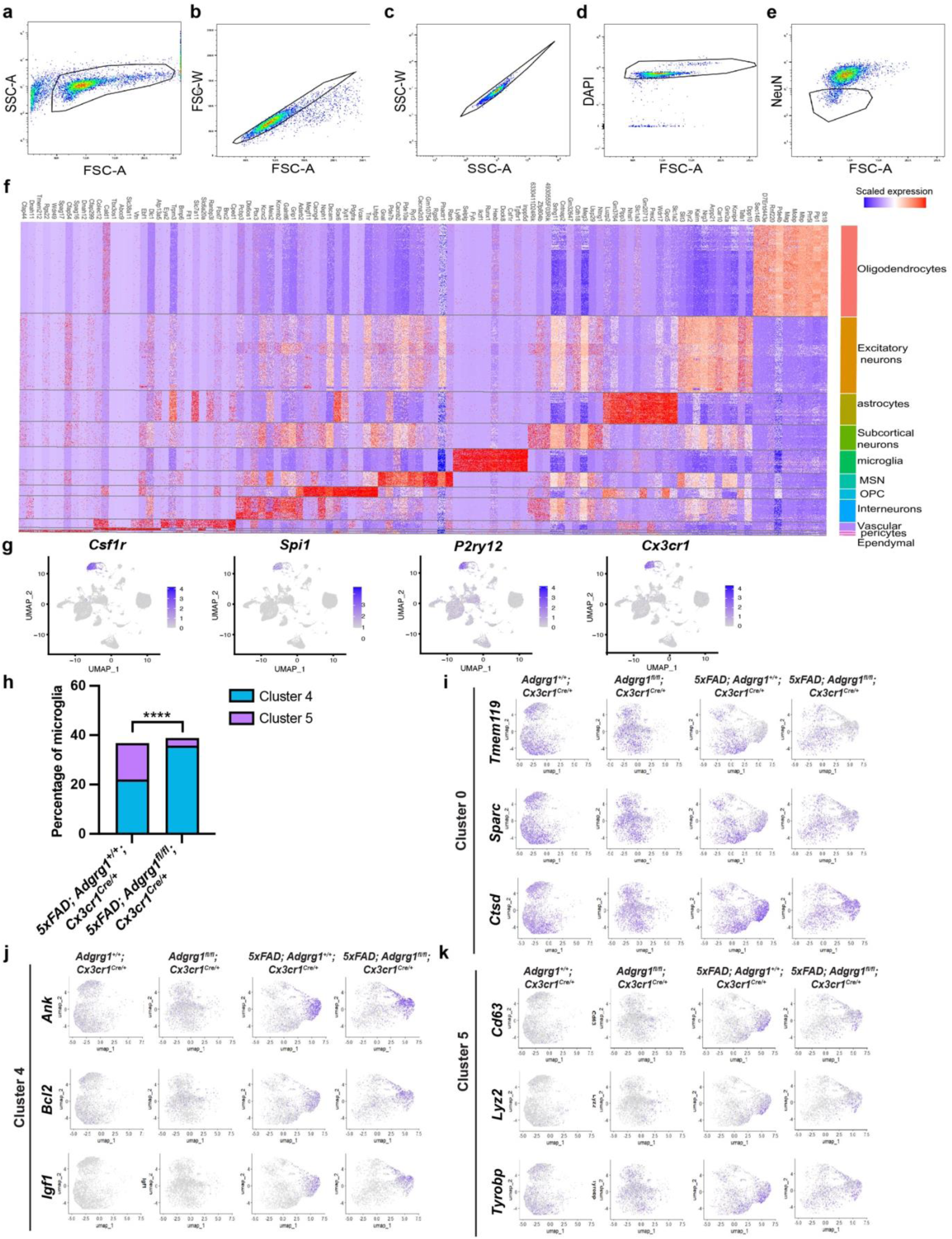
Characterization of cell-type specific markers and microglial subclusters markers in snRNA-seq, related to Figure 5. (a-e) Representative flow cytometry plots showing the gating strategy used to sort DAPI^+^ single nuclei (d), and DAPI^+^ NeuN^-^ nuclei (e). (f) Heatmap showing the expression of cell-type-specific marker genes, corresponding to identified clusters in Figure 5b. (g) Feature plots showing transcript expressions of microglia marker genes projected onto the UMAP of all cells, indicating the specificity of the microglial cluster. Color intensity indicates normalized expression levels. (h) Bar plots indicating the percentage of cells from microglia subcluster 4 and 5 between *5xFAD; Adgrg1^+/+^; Cx3cr1^Cre/+^* and *5xFAD; Adgrg1^fl/fl^; Cx3cr1^Cre/+^* mice. Chi-square test result indicates the significance of percentage difference between these two genotypes. p=6.32×10^−58^. (i-k) UMAP plots showing the gene expression enrichment for selected markers within microglial subcluster 0, 4, and 5.

**Figure S4.**
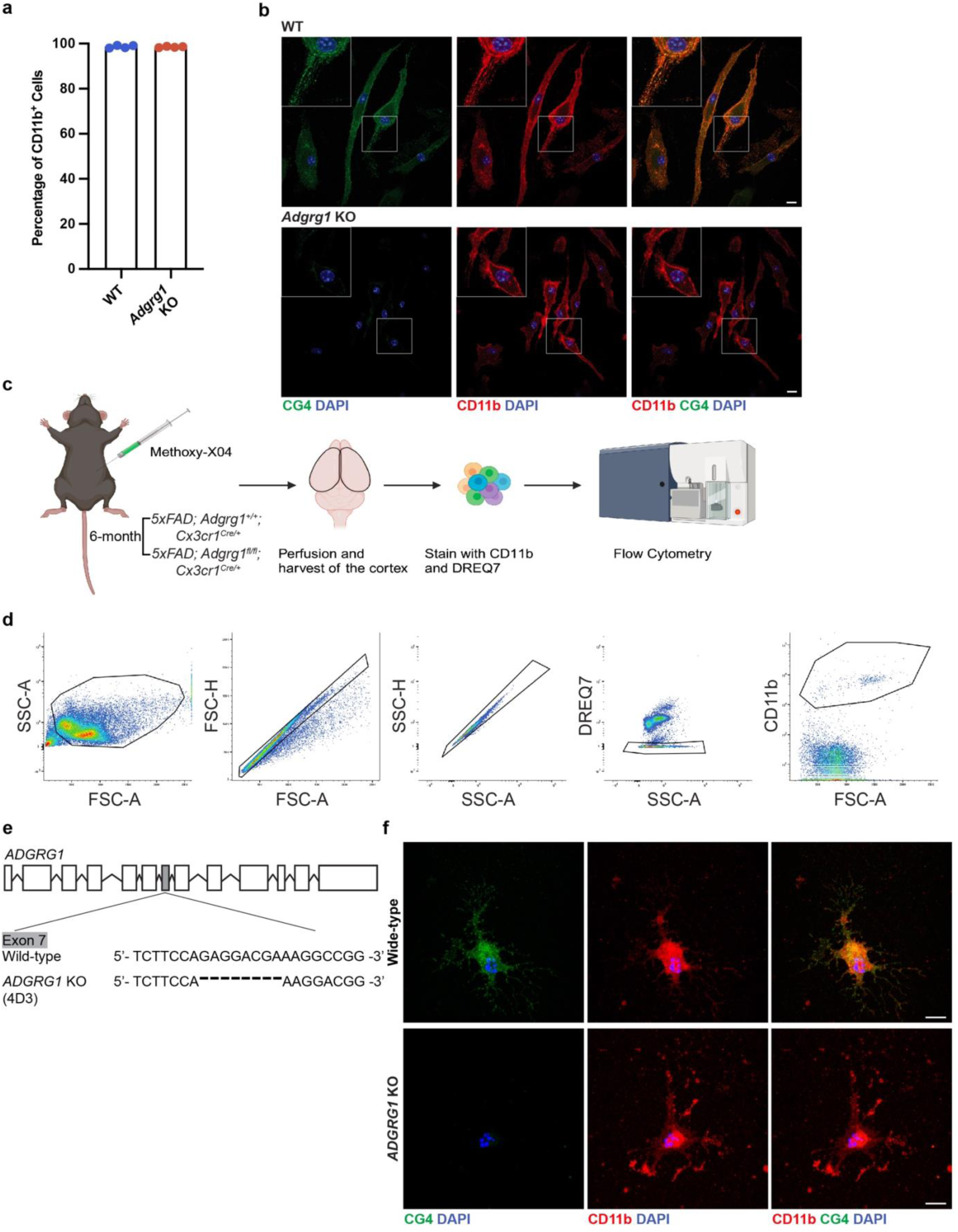
Assessment of microglial culture purity and strategy for *in vivo* Aβ engulfment assay, related to Figure 6. (a) Percentage of CD11b-positive cells relative to the total number of DAPI-positive cells. Each circle represents one biological replicate. (b) Representative immunofluorescence images of primary microglia derived from WT and *Adgrg1* KO pups, stained with CG4 (green), CD11b (red) and DAPI. Scale bar, 10 μm. (c) Schematic for the *in vivo* Aβ engulfment assay. Schematic created with BioRender. (d) Representative flow cytometry plots showing the gating strategy for analyzing microglial engulfment of Aβ *in vivo*. (e) Schematic drawing of the CRISPR/Cas9 approach to target human *ADGRG1*. (f) Representative immunofluorescence images of hESC-derived microglia from human wild-type and *ADGRG1* KO conditions, labeled with CG4 (green), CD11b (red) and DAPI. Scale bar, 10 μm.

**Figure S5.**
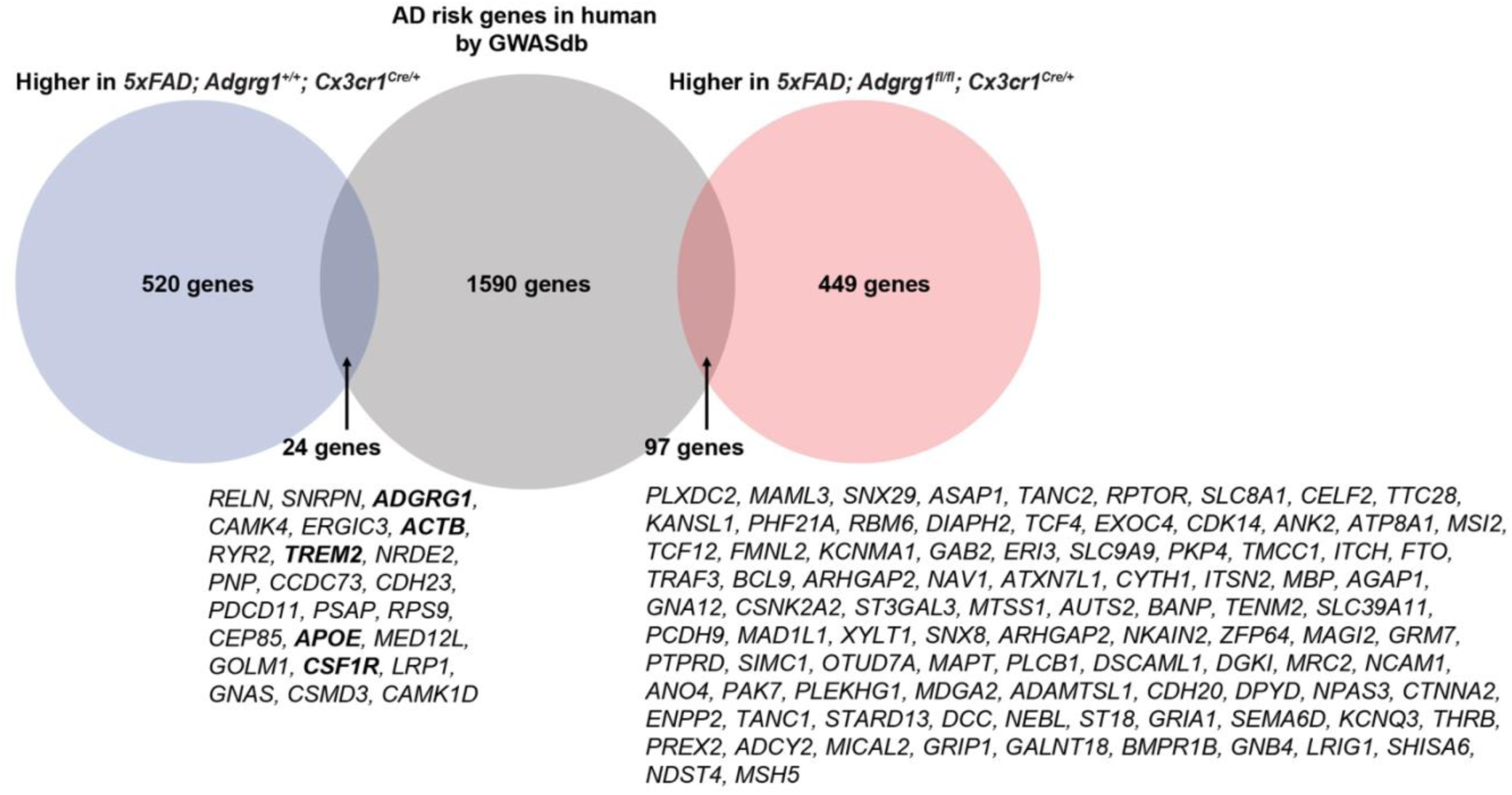
Overlap of microglial DEGs in AD mouse models with human AD risk genes, related to Figure 7. Venn diagram illustrating the overlap of DEGs in microglia from *5xFAD; Adgrg1^+/+^; Cx3cr1^Cre/+^* and *5xFAD; Adgrg1^fl/fl^; Cx3cr1^Cre/+^*mice with human AD risk genes identified in GWASdb. 24 genes were overlapped between *5xFAD; Adgrg1^+/+^; Cx3cr1^Cre/+^* microglia DEG and human AD risk genes, and 97 genes were overlapped between *5xFAD; Adgrg1^fl/fl^; Cx3cr1^Cre/+^* microglia DEG and human AD risk genes.

**Graphical abstract:**
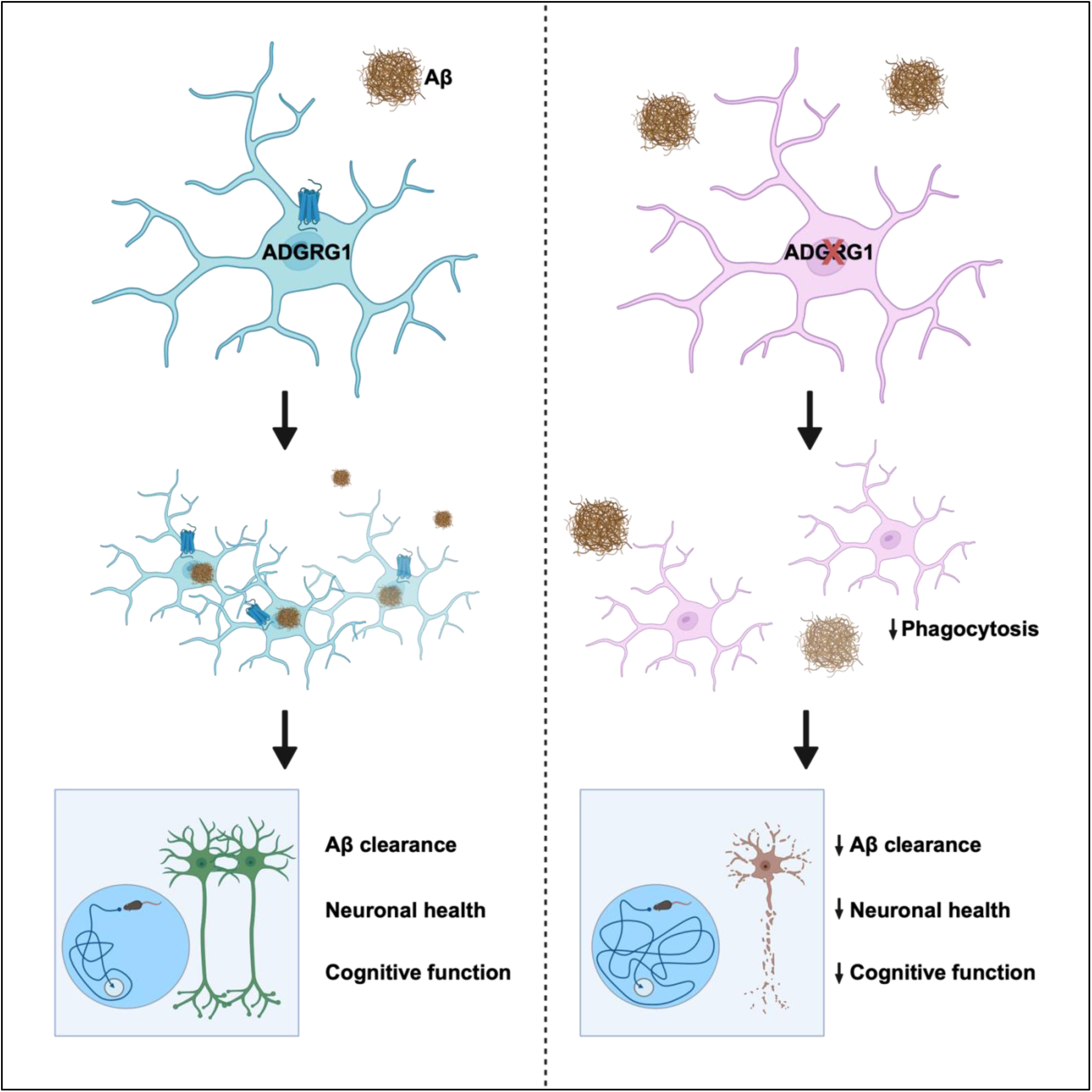
Microglial ADGRG1 regulates unique transcriptomic alterations and protective microglial responses in AD pathology. In control microglia, ADGRG1 maintains a robust homeostatic and phagocytic transcriptomic status, enhancing microglial capability to effectively clear Aβ plaques. While *Adgrg1* knockout does not prevent the transition to disease-associated microglia, it results in a distinct transcriptomic profile associated with impaired phagocytosis function. These deficiencies lead to increased Aβ deposition, exacerbate neurodegeneration, and contribute to deficits in learning and memory. Schematic created with BioRender.

**Table S1.** Clinical and Pathological Data on the Human Individuals Profiled with Immunostaining in the Study, related to Figure 7.

| Case No | Sex | Age at death | Clinical Diagnosis | Primary Neuropath Diagnosis | AD Thal Phase | AD Braak Stage | AD CERAD Neuritic Plaque Score | AD Neuropathologic change | Lewy body Disease stage | PMI (hrs) | Fixed Block Side |
| --- | --- | --- | --- | --- | --- | --- | --- | --- | --- | --- | --- |
| 1 | M | 91 | Control | None | 2 | 4 | sparse | Low | Absent | 5.6 | L |
| 2 | F | 82 | Control | None | 2 | 1 | absent | Low | Absent | 9.6 | L |
| 3 | F | 84 | Control | None | 2 | 2 | sparse | Low | Absent | 7.7 | R |
| 4 | F | 86 | Control | None | 1 | 2 | moderate | Low | Absent | 7.8 | L |
| 5 | F | 92 | Control | None | 0 | 2 | absent | Not | Absent | 8.0 | L |
| 6 | F | 86 | Control | None | 0 | 2 | absent | Not | Brainstem-predominant | 6.4 | L |
| 7 | M | 89 | MCI | ADNC | 4 | 3 | moderate | Intermediate | Absent | 9.1 | L |
| 8 | M | 77 | MCI | AGD | N/A | 2 | sparse | Low | Absent | 4.9 | L |
| 9 | M | 87 | MCI | ADNC | 1 | 4 | absent | Low | Absent | 9.7 | L |
| 10 | M | 87 | MCI | AGD | 1 | 2 | absent | Low | Absent | 6.4 | L |
| 11 | F | 88 | MCI | AGD | N/A | 3 | sparse | Low to intermediate | Absent | 15.5 | R |
| 12 | F | 81 | MCI | Low ADRC | 1 | 2 | absent | Low | Absent | 30.3 | L |
| 13 | M | 88 | Dementia | AD | 5 | 6 | frequent | High | Absent | 10.8 | L |
| 14 | M | 82 | Dementia | AD | 5 | 6 | frequent | High | Absent | 6.7 | L |
| 15 | M | 86 | Dementia | AD | 4 | 5 | frequent | High | Absent | 8.6 | L |
| 16 | M | 77 | Dementia | AD | 5 | 6 | frequent | High | Absent | 12.4 | R |
| 17 | M | 77 | Dementia | AD | 5 | 6 | frequent | High | Absent | 20.5 | L |
| 19 | M | 68 | Dementia | AD | 5 | 6 | frequent | High | Absent | 9.3 | L |
| 20 | F | 75 | Dementia | AD | 5 | 6 | frequent | High | Absent | 6.9 | R |

Alzheimer's disease (AD), AD neuropathological change (ADNC), Argyrophilic grain disease (AGD), AD-related changes (ADRC)

## METHODS

### EXPERIMENTAL MODEL AND SUBJECT DETAILS

#### Mice

All mice were handled according to the guideline of the Institutional Animal Care and Use Committee at the University of California, San Francisco (IAUCU# AN194756-01F). The *Adgrg1^fl/fl^* mice were generated in-house and have been reported previously. To confirm the presence of loxP sites, the following primers were used: (1) 5’-GGT GAC TTT GGT GTT CTG CAC GAC-3’, (2) 5’-TGG TAG CTA ACC TAC TCC AGG AGC-3’ and (3) 5’-CAC GAG ACT AGT GAG ACG TGC TAC-3’. The *Cx3cr1^Cre/+^* mice were acquired from the Jackson Laboratory (strain number: 025524). For detecting Cre expression, these primers were employed: (1) 5’-GCA GGG AAA TCT GAT GA AG-3’; (2) 5’-GAC ATT TGC CTT GCT GGA C-3’ and (3) 5’-CCT CAG TGT GAC GGA GAC AG-3’. 5xFAD mice were obtained from the Jackson Laboratory (strain number: 034848), and their genotype was determined using these primers: (1) 5’-CGG GCC TCT TCG CTA TTA C-3’; (2) 5’-ACC CCC ATG TCA GAG TTC CT-3’ and (3) 5’-TAT ACA ACC TTG GGG GAT GG-3’. The *P2ry12^CreER/+^; Ai14* mice were obtained from Dr. Thomas Arnold at UCSF. To detect *P2ry12^CreER^* expression, these primers were employed: (1) 5’-AAG AAG GTG GCG AAC CAA G-3’; (2) 5’-CAC CCTG CAG ACT AAG ATT TTT CC-3’ and (3) 5’-GTT CAG CAG GGA ACC ATT TC-3’. To induce CreER activity, one-month-old animals (postnatal day 31, P31) were intraperitoneal injected once daily with 100 mg/kg tamoxifen dissolved in corn oil at a concentration of 20 mg/mL for five consecutive days.

For our experimental design, *Adgrg1^+/+^; Cx3cr1^Cre/+^* mice were designated as Con group, *Adgrg1^fl/fl^; Cx3cr1^Cre/+^* mice as cKO mice; *5xFAD; Adgrg1^+/+^; Cx3cr1^Cre/+^* mice as 5xFAD-Con and 5xFAD; *Adgrg1^fl/fl^; Cx3cr1^Cre/+^* mice as 5xFAD-cKO. Tamoxifen-induced *Adgrg1^+/+^; P2ry12^CreER/+^; Ai14* mice as iCon; *5xFAD; Adgrg1^+/+^; P2ry12^CreER/+^; Ai14* mice as 5xFAD-iCon; 5xFAD; *Adgrg1^fl/fl^; P2ry12^CreER/+^; Ai14* mice as 5xFAD-icKO mice.

#### Generation and maintenance of WT and *ADGRG1* KO human embryonic stem cells

Experiments using hESC were conducted in accordance with UCSF guidelines and regulations. The hESC line WA09/H9 was obtained from Dr. Arnold Kriegstein’s (UCSF). All hESC were expanded on growth factor-reduced Matrigel-coated plates. hESC were grown in StemFlex Pro Media supplemented with 10 µM ROCK inhibitor Y-27632 for the first day, which was removed the following day if there were at least eight cells per colony for most colonies. Once this criterion was reached, Rock inhibitor was removed. Media were changed every other day, and cell lines were passaged when colonies reached 80% confluency. Stem cells were passaged using ACCUTASE^TM^ solution (Millipore, SCR005). All lines used for this study were below passage 30.

hESC were edited as previously described with modifications^69,70^. In brief, *ADGRG1* KO was generated using CRISPR/Cas9-based non-homology end joining, largely following the protocol of Alt-RTM CRISPR-Cas9 System from Integrated DNA Technologies (IDT). The guide RNA (gRNA) sequence (5’-ACACTCTTCCAGAGGACGAA-3’) was selected from Predesigned Alt-R CRISPR-Cas9 gRNA (IDT), targeting exon 7 of the *ADGRG1* gene locus. Equal amount of crRNA and ATTO^TM^550-labeled tracrRNA (IDT, 1075927) were mixed to a final concentration of 100 μΜ, heated to 95 °C for 5 minutes, and then cooled to room temperature to anneal, followed by forming the RNP complex with Alt-R^TM^ S.p. HiFi Cas9 Nuclease V3 (IDT,1081061) at room temperature for 20 minutes. The RNP complex was delivered to single stem cell suspension using the Neon^TM^ electroporation system (1500V, 20ms, 1 pulse) according to the manufacturer’s instructions. After electroporation, ATTO^TM^550+ cells were selected by FACS after three days of culture and sparsely seeded to form single-cell colonies. A loss-of-function mutation cell line 4D3 was selected by Sanger sequencing, followed by exclusion of any mutations at the top 5 potential off-target sites. Further Sanger sequencing and immunostaining were applied to confirm *ADGRG1* KO.

#### Generation of hESC-derived microglia

Cells were differentiated into iMG following the protocols of the STEMdiff^TM^ Hematopoietic Kit (05310, STEMCELL), STEMdiff^TM^ Microglia Differentiation Kit (100-0019, STEMCELL), and STEMdiff^TM^ Microglia Maturation Kit (100-0020, STEMCELL). On day 28 of maturation, cells were collected and seeded at a density of 20,000 cells per well in PLL-coated 96-well plates and 80,000 cells on PLL-coated coverslips in 24-well plates for further experiments and maintained in the microglia maturation medium.

#### Purification of primary microglia

Primary microglia were isolated from mixed glial cultures derived from the cortices of postnatal day 3 to 5 (P3-P5) 5xFAD; *Adgrg1^fl/fl^; Cx3cr1-Cre^+/-^* and *5xFAD; Adgrg1^+/+^; Cx3cr1-Cre^+/-^* mice. The cortical tissues were dissected in Hanks’ Buffer Saline Solution (HBSS) and the meninges were carefully removed. Tissues were finely minced on ice using a blade and then transferred into 3 mL of HBSS. After allowing the tissue fragments to settle at the bottom of the tube, the supernatant was gently removed. The resulting cell pellets were transferred in 0.01% poly-L-Lysine (PLL)-coated T75 flasks containing 10 mL of mixed glial culture medium composed of high-glucose DMEM (Cytiva, SH30243.01), supplemented with 20% heat-inactivated fetal bovine serum (FBS; Thermo Fisher, A31604-02), 1% Pen/strep (Thermo Fisher, 15070063), and 20 µg/mL GM-CSF (PeproTech, 315-03). On the third day *in vitro* (DIV3), the culture medium was completely replaced to remove non-adherent cells and debris. On DIV10, microglia were flushed off the astrocyte layer by shaking at 200 rpm for 2 hours within a standard tissue culture incubator. Subsequently, the detached microglia were collected and centrifuged at 800 rpm for 5 minutes at 4°C. The purified microglia were then seeded at a density of 20,000 cells per well in PLL-coated 96-well plates and 80,000 cells on PLL-coated coverslips in 24-well plates for further experiments. The microglia were cultured in microglia culture medium consisting of DMEM/F12 (Millipore Sigma, D8437), supplemented with 1% Pen/strep, 2mM L-glutamine (Thermo Fisher, 25030081), and 5 µg/mL N-Acetyl Cysteine (NAC; Millipore Sigma, A7250), 5 µg/mL Insulin (Millipore Sigma, I0516), 100 µg/mL Apo-transferrin (Millipore Sigma, T2252), 100 ng/mL Sodium Selenite (Millipore Sigma, S5261), 2 ng/mL TGF-beta (PeproTech, 100-35B), 100 ng/mL IL-34 (R&D Systems, 5195) and 1.5 µg/mL Cholesterol (Millipore sigma, C3045). Microglia were incubated overnight in this medium before being subjected to subsequent treatments.

#### Human brain tissue samples

Postmortem human brain samples were obtained from the Neurodegenerative Disease Brain Bank at the University of California, San Francisco, under approved ethical guidelines. Formalin-fixed paraffin-embedded sections from middle temporal gyrus from seven individuals diagnosed with AD, five individuals with MCI, and six neurologically healthy control subjects were analyzed by immunostaining. Detailed information regarding the age, sex, postmortem interval (PMI), and other relevant clinical data of the tissue donors are documented in **Table S1**.

### METHOD DETAILS

#### Mouse brain sample preparation

The brain samples were collected from mice following a standardized and ethically approved procedure. The mice were anesthetized using an isoflurane chamber and subsequently underwent transcardial perfusion with 20 mL of ice-cold 1x PBS. The right hemisphere of each brain was dissected into cortex, hippocampus, mid-brain and cerebellum. These tissues were immediately snap-frozen in dry ice and then stored at -80°C. The left hemisphere was fixed in 4% paraformaldehyde (PFA; Thermo Fisher, J19943-K2) for 48 hours at 4°C. Following fixation, the tissue was cryoprotected by immersion in 30% sucrose in PBS until it sank. The cryoprotected tissue was then embedded in O.C.T. (SCIgen, #4586). The embedded brain tissue was sectioned at thicknesses of either 14 µm or 40 µm using a cryostat (Leica) for the subsequent immunohistochemistry.

#### Immunofluorescence staining

For the 14-µm brain sections, antigen retrieval was performed for 5 minutes at 95°C using an antigen retrieval buffer (BD Pharmingen, 550524). Subsequently, sections were blocked in a blocking buffer containing 5% goat serum and 1% bovine serum albumin (BSA), and 0.03% Triton X-100 in PBS at room temperature for 1 hour. Slices were incubated with primary antibodies overnight at 4°C. The antibodies used were as follows: For amyloid plaque labeling, anti-Aβ (MOAB2, 1:1000; or H31L21, 1:1000). To quantify microglial cells, anti-Iba1 (1:500). For neuronal health assessment, anti-NeuN (1:500), anti-CTIP2 (1:1000), anti-TBR1 (1:1000), anti-MAP2 (1:2000), anti-LAMP1 (1:200), anti-PSD95 (1:500), anti-vGlut2 (1:500). For enhanced permeability in anti-ADGRG1 (CG4, 1:200) staining, sections underwent a 15-minute treatment with Protease III (ACD) at 40°C before blocking. The following secondary antibodies were applied at a 1:500 dilution at room temperature for 2 hours: goat anti-chicken IgY Alexa Flour 555, goat anti-rat IgG Alexa Flour 555, goat anti-mouse IgG Alexa Flour 488, goat anti-mouse IgG Alexa Flour 647, goat anti-rabbit IgG Alexa Flour 488, goat anti-rabbit IgG Alexa Flour 647, goat anti-guinea pig IgG Alexa Flour 488, goat anti-guinea pig IgG Alexa Flour 555. DAPI (1 µg/mL) was used for nuclear counterstaining, followed by mounting with Fluoromount-G mounting medium. For human brain sections, to reduce autofluorescence, slides were treated with TrueBlack Lipofuscin Autofluorescence Quencher at a 1:20 dilution in 70% ethanol for 30 seconds prior to mounting. Additionally, Thioflavin S staining was performed post-antibody labeling. Briefly, slides were stained with 1% Thioflavin S for 10 minutes, followed by washing in 70% ethanol for 5 minutes once and twice in water for 5 minutes. For analyzing of microglial engulfment of Aβ, 40µm free-floating sections were used. Post-incubation in 10% Triton X-100 in PBS for 30 minutes and subsequent blocking, sections were incubated with primary antibodies (anti-Aβ, anti-Iba1, anti-CD68) for 40-48 hours at 4°C and secondary antibodies were incubated for 2 hours at room temperature. Quantitative analysis of Aβ engulfment by microglia or CD68 co-localization was conducted using 3D rendering of confocal images in Imaris 9.8.0 software.

For immunocytochemistry, primary microglia seeded on coverslips were treated with Protease III at 40°C for 15 minutes for CG4 staining. Post-blocking, cells were incubated with anti-CD11b (1:200) and anti-CG4 (1:200) overnight at 4°C. Secondary antibodies were applied at a 1:500 ratio and incubated at room temperature for 1 hour. Coverslips were mounted using DAPI-Fluoromount-G.

Both mouse and human sections were imaged using a Keyence BZ-X800 microscope with a 20x objective for whole section scans and Leica SP5 confocal microscope with 20x or 63x objectives. The imaging resolution was set to 2048 x 2048 pixels, resulting in a pixel size of 550 µm x 550 µm for 20x and 174.60 µm x 174.60 µm for 63x. Z-stacks of 10 µm were acquired with a 0.5 µm z-step size, using sequential scans with 3x averaging at 488, 555, and 647 nm wavelengths and 1x averaging at 405 nm wavelength. Experiments were blinded to the genotype during image acquisition and processing.

#### RNAscope *in situ* hybridization

The RNAscope *in situ* hybridization was employed on fixed, frozen mouse, and human brain tissue samples using the Multiplex Fluorescence v2 kit (Advanced Cell Diagnostics) according to the manufacturer’s protocol. Probes for Human-*ADGRG1* and mouse-*Gpr56* were commercially available from the manufacturer. TrueBlack Lipofuscin Autofluorescence Quencher was applied to the sections prior to mounting.

#### *In vivo* Aβ phagocytosis assay

*In vivo* Aβ phagocytosis assay was adapted from a previous study with modification.^71^ 6-month-old 5xFAD; *Adgrg1^fl/fl^; Cx3cr1-Cre^+/-^* and *5xFAD; Adgrg1^+/+^; Cx3cr1-Cre^+/-^* mice were intraperitoneally injected with 10 mg/kg of methoxy-X04 in a solution of 10% DMSO, 45% propylene glycol, and 45% PBS. Three hours later, mice were anesthetized using an isoflurane chamber and transcardially perfused with 20 mL of ice-cold 1x PBS. The cortices, dissected on ice, were homogenized in 1 mL of 1xPBS, then centrifuged at 800g for 5 minutes at 4°C. The resulting cell pellet was resuspended in 10 mL of 30% Percoll and overlaid with 2 mL of 1x PBS. This was centrifuged at 800g for 15 minutes at 4°C to separate myelin. After removing myelin, cells were washed in 1 mL of FACS buffer (1% BSA in PBS) and pelleted at 300g for 5 minutes at 4°C. The cells were blocked with FACS buffer containing CD16/CD32 (1:100) for 20 minutes on ice, then incubated with DRAQ7 (1:200), and CD11b-PE (1:50) in FACS buffer for 20 minutes on ice. After a final centrifugation, cells were resuspended in FACS buffer and analyzed by flow cytometry to gate CD11b^+^ microglia and assess methoxy-X04 signal within the gated microglia population.

#### pHrodo-oligomeric Aβ preparation

Oligomeric Aβ was prepared following a previously published protocol.^72^ Briefly, HFIP pre-treated Aβ was dissolved in DMSO and further diluted in phenol-red free F-12 medium to obtain a final concentration of 100 µM. The samples were sonicated for 10 minutes, followed by a 24-hour incubation at 4°C. The samples were centrifuged at 14,000g for 10 minutes. The resultant supernatant was aliquoted and stored at -80°C. Oligomeric Aβ was then incubated with pHrodo Red-Succinimidyl Ester diluted in 0.1M sodium bicarbonate for 30 minutes covered at room temperature. Following incubation, HBSS was used to wash the samples and remove excess dye. pHrodo Red-labelled Aβ was used immediately in subsequent experiments.

#### *In vitro* phagocytosis assay

Microglial cells, both mouse primary microglia and hESC-derived microglia, were seeded in clear bottom black 96-well plates (Corning, 07-200-565). 24 hours later after seeding, the cells were treated with 1 µM of pHrodo Red-labeled oligomeric Aβ. Following the treatment, microglia were immediately subjected to live-cell imaging using the Incucyte S3 Live-Cell Analysis System (Essen Bioscience). Phase contrast and RFP images were captured for each well at hourly intervals over a 24-hour period. For each time point, the integrated intensity of red objects was quantified. This quantification was performed using Red Calibrated Unit (RCU) measurements, multiplied by the area (µm^2^) of recorded image.

#### General design of behavioral tests

Behavioral tests were conducted on 4-month-old male and female mice, housed in groups of two to five mice per cage. All procedures were carried out at the UCSF Gladstone Behavioral Core by professional technicians. The genotypes of all mice were blinded to both experiment conductors and data analyzers.

##### Rotarod

The rotarod test was performed on a rotarod apparatus (Med Associates Inc., Vermont, USA) under normal lighting conditions. The rotarod apparatus was cleaned with Vimoba before and after each use, and sanitized with 70% alcohol between individual trials. During training sessions, groups of up to five mice, segregated by sex, were simultaneously placed on the rotarod. The rotarod was set to rotate at a constant speed of 16 rotations per minute (rpm). A trial was deemed complete when a mouse fell off the rod or after a duration of 5 minutes had elapsed. Each mouse underwent 3 individual trials with an inter-trial interval of 15**-**20 minutes. For the actual test, the rotarod speed was progressively increased from 4 rpm to 40 rpm, with an increment of 4 rpm every 30 seconds. The mice were subjected to two sessions of three trials each, conducted in the morning and afternoon over two consecutive days.

##### Morris water maze

The Morris water maze consisted of a circular pool (122 cm diameter, 50 cm height) filled with water maintained at 22± 1°C. The water was made opaque using non-toxic white tempera liquid paint to obscure the submerged platform. Mice underwent hidden platform training over 5 days, with 2 sessions comprising 2 trials each day. The submerged platform (1.5 cm x 1.5 cm) was kept in a consistent location throughout the training period, located 1.5 cm below the water surface and without visible cues. Mice were given a maximum of 60 seconds to locate the platform in each trial and were required to stay on the platform for at least 10-15 seconds before being removed. The starting positions for each trial were randomized, including both "closer" and "further" drop locations relative to the platform. 24 hours following the final day of training, probe trials were conducted with the platform removed. The mice’s swim patterns were recorded for 60 seconds using the Etho VisionXT video tracking system (Noldus, Netherlands). For each trial, mice were introduced into the pool at a position 180° opposite the former platform location. After the 60-second trial, the experimenter places his/her hand where the hidden platform used to be and lets the mouse swim to his/her hand.

#### Isolation of nuclei from fresh frozen mouse tissues

Flash-frozen mouse neocortex brain tissues were minced on dry ice and transferred to a Dounce homogenizer. The tissues were homogenized in 1.5 mL of Nuclei-Pure Lysis Buffer (Sigma) supplemented with 0.2 U/μL RNase inhibitor (Takara) on ice. The homogenate was filtered through a 40 µm cell strainer to remove large debris and centrifuged at 600g for 5 minutes at 4°C to pellet the nuclei. The pelleted nuclei were washed thrice with 1 mL of nuclei wash buffer, composed of 1% BSA in PBS, 20 mM DTT, and RNase inhibitor, with centrifugation between each wash. The washed nuclei were resuspended in 0.04% BSA in PBS containing RNase inhibitor and incubated with anti-NeuN-Alexa 488 conjugated antibody (1:500) for 40 minutes on ice. Subsequently, the nuclei were stained with DAPI (1:100) for 5 minutes on ice. Nuclei were washed and passed through a 35 µm cell strainer cap of Falcon round-bottom polystyrene test tubes. Using flow cytometry, DAPI^+^ and DAPI^+^ NeuN^-^ nuclei were gated and collected from each animal. The nuclei were then pelleted again at 600g for 5 minutes at 4°C and resuspended in 0.04% BSA in PBS. The nuclei were counted for downstream library preparation procedures.

#### Single-nucleus RNA sequencing

Following nuclei isolation, the snRNA-seq library was prepared using Chromium Next GEM Single Cell 3′ Reagent Kits v3.1 according to the manufacturer’s protocol (10X Genomics). The generated snRNA-seq libraries were sequenced using NovaSeq 6000 aiming for 50K read pairs per nuclei. Sequencing results were aligned to the mm10 mouse genome (GENCODE vM23/Ensembl 98) using CellRanger v6.1.2 (10x Genomics). Include-introns was used to include pre-mature mRNA in the nucleus. The count matrix then underwent doublet removal step using DoubltFinder v2.0.3 (https://www.cell.com/cell-systems/fulltext/S2405-4712(19)30073-0) accompanied with Seurat v4.1.1 (https://www.sciencedirect.com/science/article/pii/S0092867421005833?via%3Dihub). Nuclei with 500-6000 genes detected, 1000-40,000 unique molecular identifiers (UMIs), and less than 2% reads mapped to the mitochondrial genes were kept for further analysis. Clustering and cell-types markers were identified using Seurat v4.1.1.

#### Clustering and finding markers

Principal component analysis was performed prior to clustering, and number of principal components used were determined based on the ElbowPlot. For initial clustering of all cell-types, 20 principle components were used. And 7 were used for subclustering of microglia. Clustering was performed using the FindClusters function, which works on K-nearest neighbor (KNN) graph model with the granularity ranging from 0.1–0.9 and selected 0.4 for the downstream clustering. We performed differential expression of each cluster against all other clusters, to identify negative and positive markers for that cluster. Nuclei from microglia were taken and re-clustered to further analyze the subclusters.

#### Analysis of gene differential expression

Differential expression of genes between conditions was done using MATLAB. p-value were calculated using Student’s t-test, accompanied with Storey’s multi-test correction. Genes with at least 33% change in the expression level and q-value < 0.05 are used in gene ontology (GO) analysis using Enrichr.

#### Projecting Trem2 dataset to microglia subcluster UMAP

To compare microglia subclustering in the *Trem2* dataset, we projected all microglia (3713 cells) from the Trem2 dataset on to our microglial subclustering using ProjectUMAP function in Seurat.

#### Chi-square analysis

Chi-square test of independence was employed to determine where there was significant association between cell populations in cluster 4 and 5 under the two experimental conditions *5xFAD; Adgrg1^+/+^; Cx3cr1^Cre/+^* and *5xFAD; Adgrg1^fl/fl^; Cx3cr1^Cre/+^*.

#### Human ADGRG1 expression correlation analysis

To perform transcriptomic meta-analysis, original count matrix of 590 human brain samples were collected from published dataset.^50–54^ These microglial transcriptomes are pooled by sample, and Pearson’s correlation coefficient is calculated between the expression level of all genes and *ADGRG1*. The correlation coefficient is then plotted as a histogram to show the distribution. Important genes in the proposed pathway are labeled along with the number of genes in that bin on the histogram.

## QUANTIFICATION AND STATISTICAL ANALYSIS

Mean values, SEM values, Student’s *t* test, one-way ANOVA, and two-way ANOVA were calculated using Prism 10.0 software (GraphPad). *p* value less than 0.05 were considered significant. ns=not significant, *p < 0.05, ** p< 0.01, *** p<0.001, ****p< 0.0001.

## RESOURCE AVAILABILITY

### Data and code availability

Original single-nuclei RNA-seq data is deposited at GEO: GSE273690 and will be publicly available as of the date of publication. Original code or any additional information required to reanalyze the data reported in this paper will be shared by the lead contact upon request.

### Lead contact

Further information and requests for resources and reagents should be directed to and will be fulfilled by the lead contact, Xianhua Piao.

## Notes

### Competing Interest Statement

The authors have declared no competing interest.

